# Unlocking the hidden genetic diversity of varicosaviruses, the neglected plant rhabdoviruses

**DOI:** 10.1101/2022.09.19.508500

**Authors:** Nicolás Bejerman, Ralf G. Dietzgen, Humberto Debat

**Affiliations:** Instituto de Patología Vegetal – Centro de Investigaciones Agropecuarias – Instituto Nacional de Tecnología Agropecuaria (IPAVE-CIAP-INTA), Camino 60 Cuadras Km 5,5 (X5020ICA), Córdoba, Argentina; Consejo Nacional de Investigaciones Científicas y Técnicas. Unidad de Fitopatología y Modelización Agrícola, Camino 60 Cuadras Km 5,5 (X5020ICA), Córdoba, Argentina; Queensland Alliance for Agriculture and Food Innovation, The University of Queensland, St. Lucia, Queensland 4072, Australia

**Author notes:** Corresponding authors: Nicolás Bejerman,; Dietzgen Ralf,; Debat Humberto.

**Keywords:** plant rhabdovirus, varicosaviruses, genome architecture, virus taxonomy, metatranscriptomics

## Abstract

The genus *Varicosavirus* is one of six genera of plant-infecting rhabdoviruses. Varicosaviruses have nonenveloped flexuous rod-shaped virions and a negative-sense, single-stranded RNA genome. A distinguishing feature of varicosaviruses, that is shared with dichorhaviruses, is a bi-segmented genome. Before 2017, a sole varicosavirus was known and characterized, then two more varicosaviruses were identified through high-throughput sequencing in 2017 and 2018. More recently, the number of known varicosaviruses has substantially increased in concert with the extensive use of high-throughput sequencing platforms and data mining approaches. The novel varicosaviruses revealed not only sequence diversity but also plasticity in terms of genome architecture, including a virus with a tentatively unsegmented genome. Here, we report the discovery of 45 novel varicosavirus genomes, which were identified in publicly available metatranscriptomic data. Identification, assembly, and curation of raw Sequence Read Archive reads resulted in 39 viral genome sequences with full-length coding regions and 6 with nearly complete coding regions. Highlights of the obtained sequences include eight varicosaviruses with unsegmented genomes, linked to a phylogenetic clade associated with gymnosperms. These findings resulted in the most complete phylogeny of varicosaviruses to date and shed new light on the phylogenetic relationships and evolutionary landscape of this group of plant rhabdoviruses. Thus, the extensive use of sequence data mining for virus discovery has allowed unlocking of the hidden genetic diversity of varicosaviruses, the largely neglected plant rhabdoviruses.

## 1. Introduction

The recently discovered huge number of diverse viruses has revealed the complexities of the evolutionary landscape of replicating entities and the challenges associated with their classification [1], leading to the first comprehensive proposal of the virus world megataxonomy [2]. Nevertheless, a minuscule portion, probably a small fraction of one percent of the virosphere has been characterized so far [3]. Therefore, we have a limited knowledge of the vast world virome, with its remarkable diversity and including every potential host organism assessed so far [4–6]. Data mining of publically available transcriptome datasets has become an efficient and inexpensive strategy to unlock the diversity of the plant virosphere [5]. Data-driven virus discovery relies on the vast number of available datasets on the Sequence Read Archive (SRA) of the National Center for Biotechnology Information (NCBI). This resource, which is growing at an exceptional rate and includes data of a large and diverse number of organisms, represents a substantial fraction of species that populate our planet, which makes the SRA database an invaluable source to identify novel viruses [7].

*Varicosavirus* is one of the six genera that are comprised of plant rhabdoviruses (family *Rhabdoviridae*, subfamily *Betarhabdovirinae*), and its members are thought to have a negative-sense, single-stranded, bisegmented RNA genome [8]. Nevertheless, recently we described the first apparently unsegmented varicosavirus [9]. In those varicosaviruses with segmented genomes, RNA 1 consists of one to two genes, with one of those encoding the RNA-dependent RNA polymerase L, while RNA 2 consists of three to five genes, with the first open reading frame (ORF) encoding a nucleocapsid protein (N) [8, 10]. On the other hand, the only unsegmented varicosavirus described so far has 5 ORFs, in the order 3’-N-Protein 2-Protein 3– Protein 4–L-5’ [9]. Varicosaviruses appear to have a diverse host range including dicots, monocots, gymnosperms, ferns, and liverworts [6, 9]. The vector of a sole member, lettuce big vein-associated virus (LBVaV) has been characterized, which is the chytrid fungus *Olpidium spp*. [11].

Until 2017, LBVaV was the only identified and extensively characterized varicosavirus [12–14], then, in 2017 and 2018, two novel varicosaviruses were identified through high-throughput sequencing (HTS) [15–16]. However, in 2021 and 2022, there was a five-fold increase in the number of reported varicosaviruses, with 12 out 15 discovered through data mining of publically available transcriptome datasets [6, 9, 17–18], while the other three were identified using HTS [19–21] (Suppl. Fig. 1).

Nevertheless, only some minor biological aspects, such as mechanical transmissibility, of few of these members were further characterized [15, 20]. Therefore, varicosaviruses are, by far, the least studied plant rhabdoviruses, and many aspects of their epidemiology remain elusive. In terms of genetic diversity, before this study, while greatly expanded by recent works, the *Varicosavirus* genus includes only three accepted species and 15 tentative members.

In this study, we identified 45 novel varicosaviruses by analyzing publicly available metatranscriptomic data. Thus, the extensive use of data mining for virus discovery has allowed us to unlock some of the hidden diversity of varicosaviruses, the much neglected plant rhabdoviruses.

## 2. Material and Methods

### 2.1 Identification of plant rhabdovirus sequences from public plant RNA-seq datasets

Three strategies were used to detect varicosavirus sequences: 1) amino acid sequences corresponding to the nucleocapsid and polymerase proteins of known varicosaviruses were used as query in tBlastn searches with parameters word size = 6, expected threshold = 10, and scoring matrix = BLOSUM62, against the Viridiplantae (taxid:33090) Transcriptome Shotgun Assembly (TSA) sequence databases. The obtained hits were manually explored and based on percentage identity, query coverage and E-value (>1e-5), shortlisted as likely corresponding to novel virus transcripts, which were then further analyzed. 2) Raw sequence data corresponding to the SRA database associated with the 1K study [22] was explored for varicosa-like virus sequences. 3) the Serratus database was explored, employing the serratus explorer tool [5] using as query the sequences of LBVaV, red clover varicosavirus and black grass varicosavirus. Those SRA libraries that matched the query sequences (alignment identity > 45%; score > 10) were futher explored in detail.

### 2.2 Sequence assembly and identification

The nucleotide (nt) raw sequence reads from each SRA experiment, which are associated with different NCBI Bioprojects (Table 1), were downloaded, and pre-processed by trimming and filtering with the Trimmomatic tool as implemented in http://www.usadellab.org/cms/?page=trimmomatic. The resulting reads were assembled *de novo* with rnaSPAdes using standard parameters on the Galaxy.org server. The transcripts obtained from *de novo* transcriptome assembly were subjected to bulk local BLASTX searches (E-value < 1e^-5^) against a collection of varicosavirus protein sequences available at https://www.ncbi.nlm.nih.gov/protein?term=txid140295[Organism]. The resulting viral sequence hits of each bioproject were visually explored. Tentative virus-like contigs were curated (extended or confirmed) by iterative mapping of each SRA library’s filtered reads. This strategy used BLAST/nhmmer to extract a subset of reads related to the query contig, used the retrieved reads to extend the contig and then repeated the process iteratively using as query the extended sequence. The extended and polished transcripts were reassembled using Geneious v8.1.9 (Biomatters Ltd.) alignment tool with high sensitivity parameters.

**Table 1.**
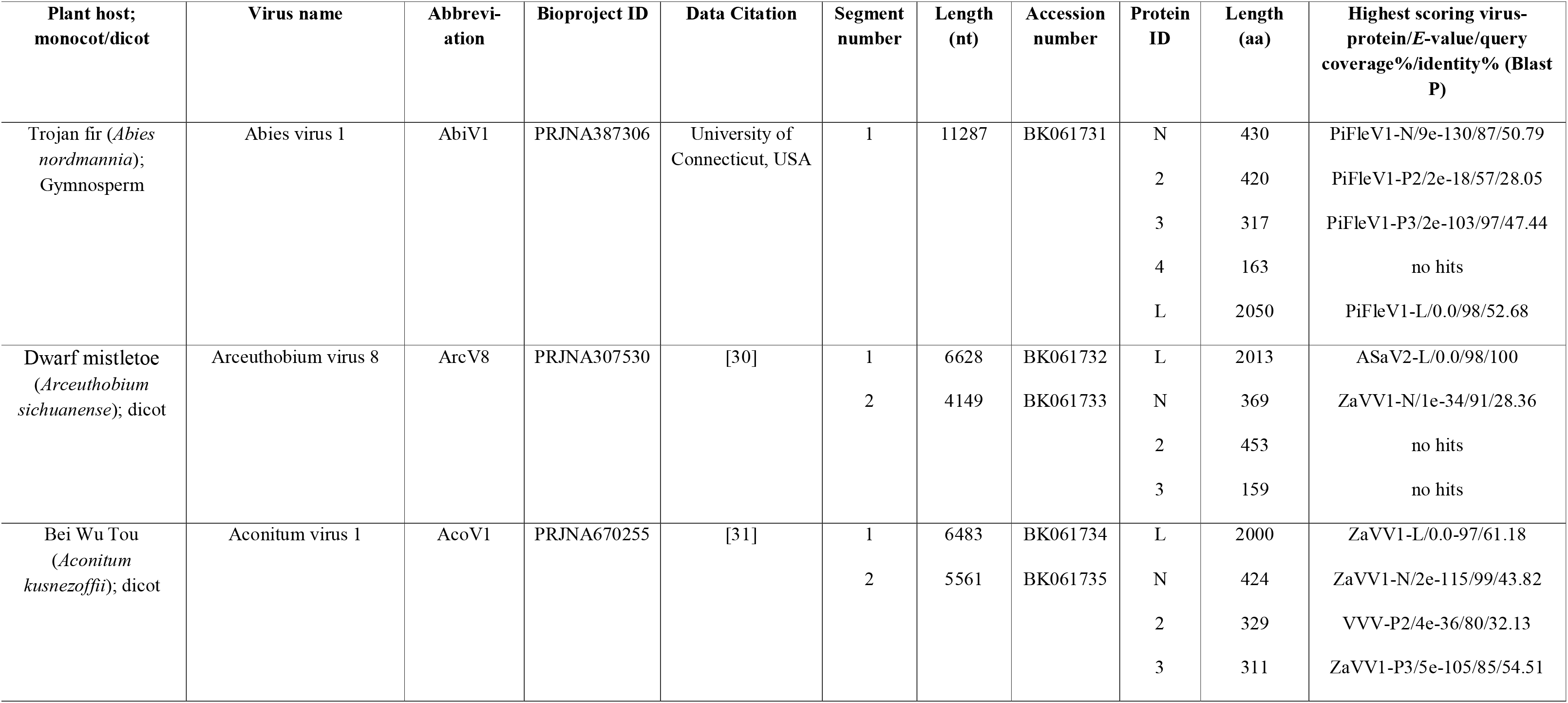
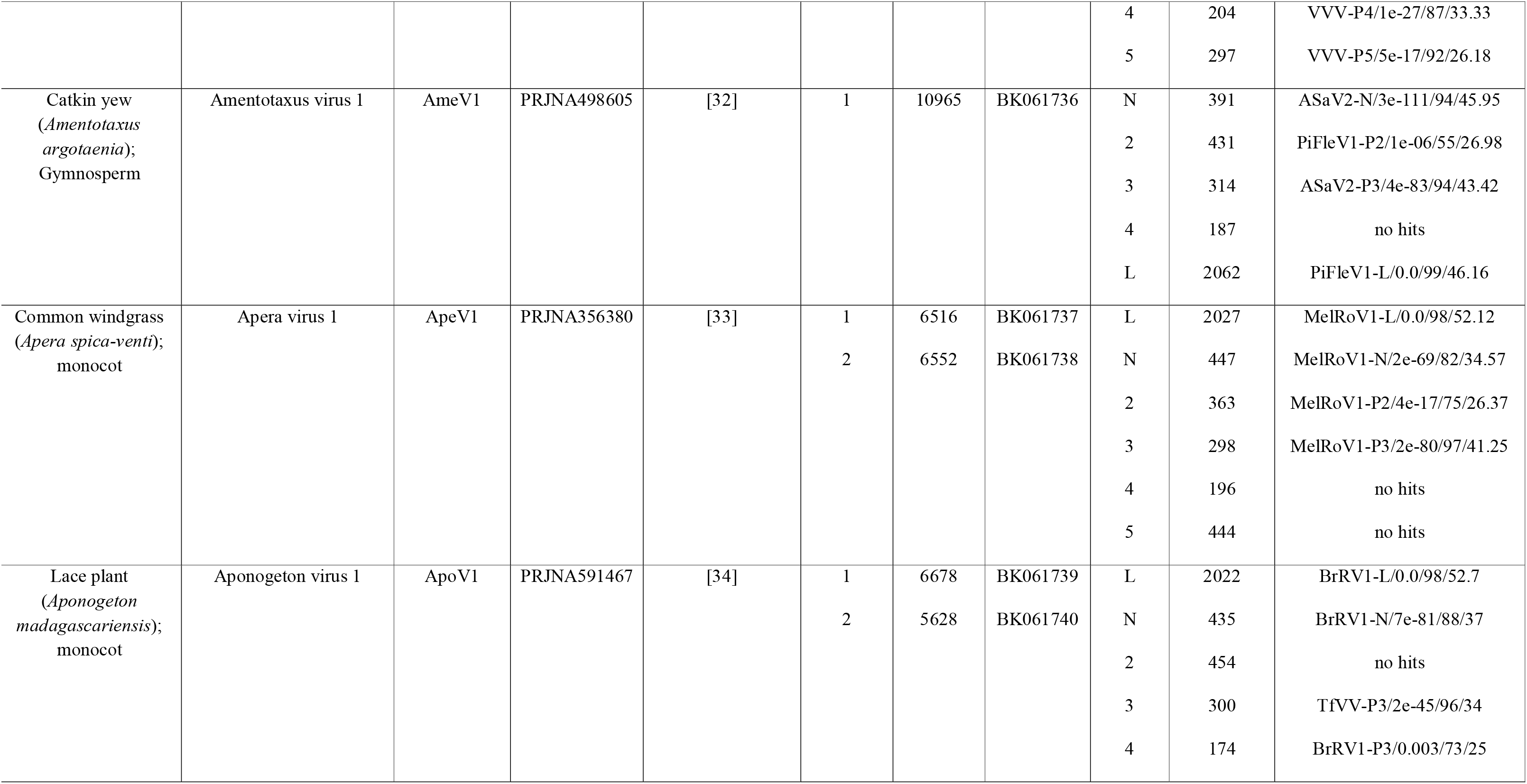
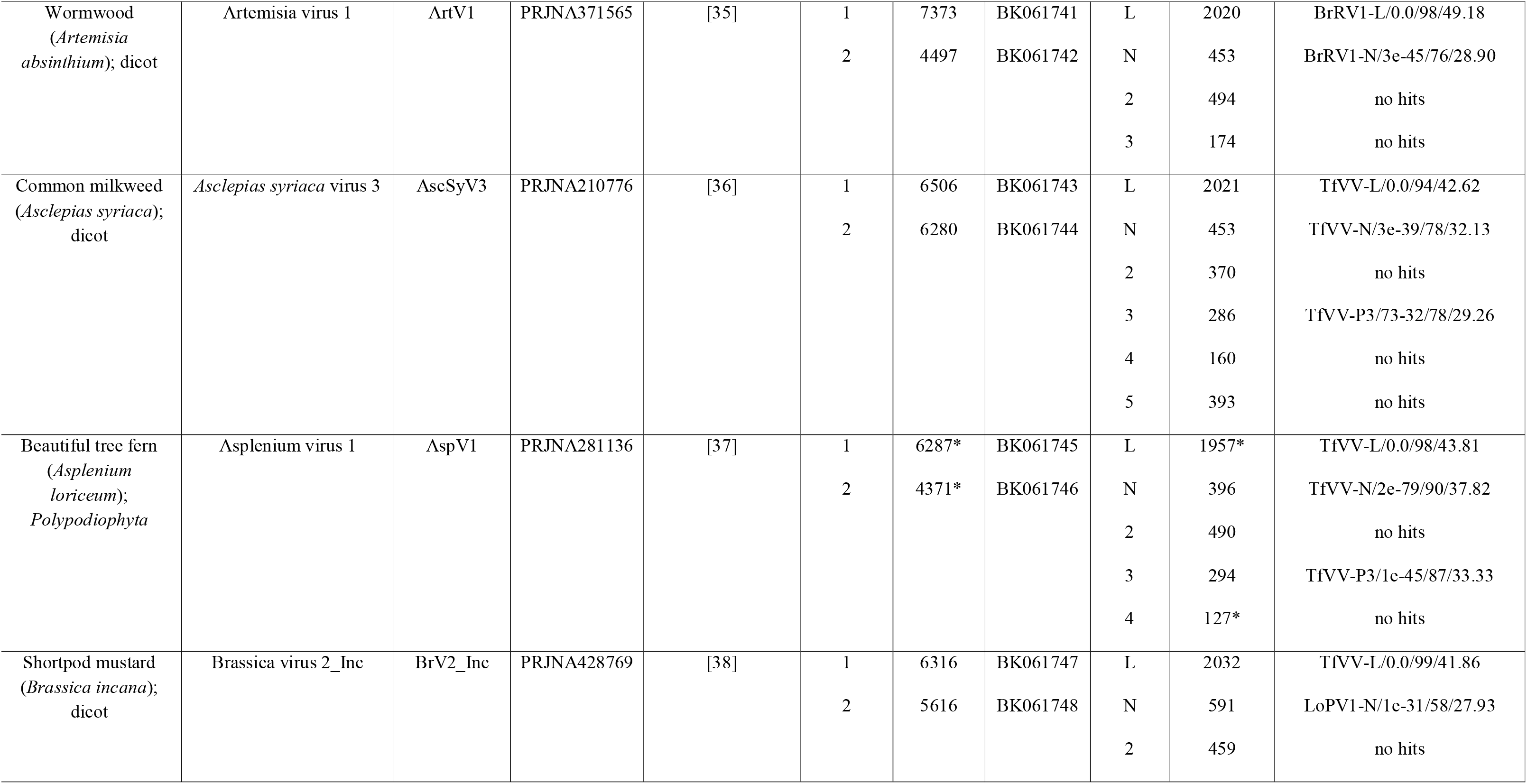
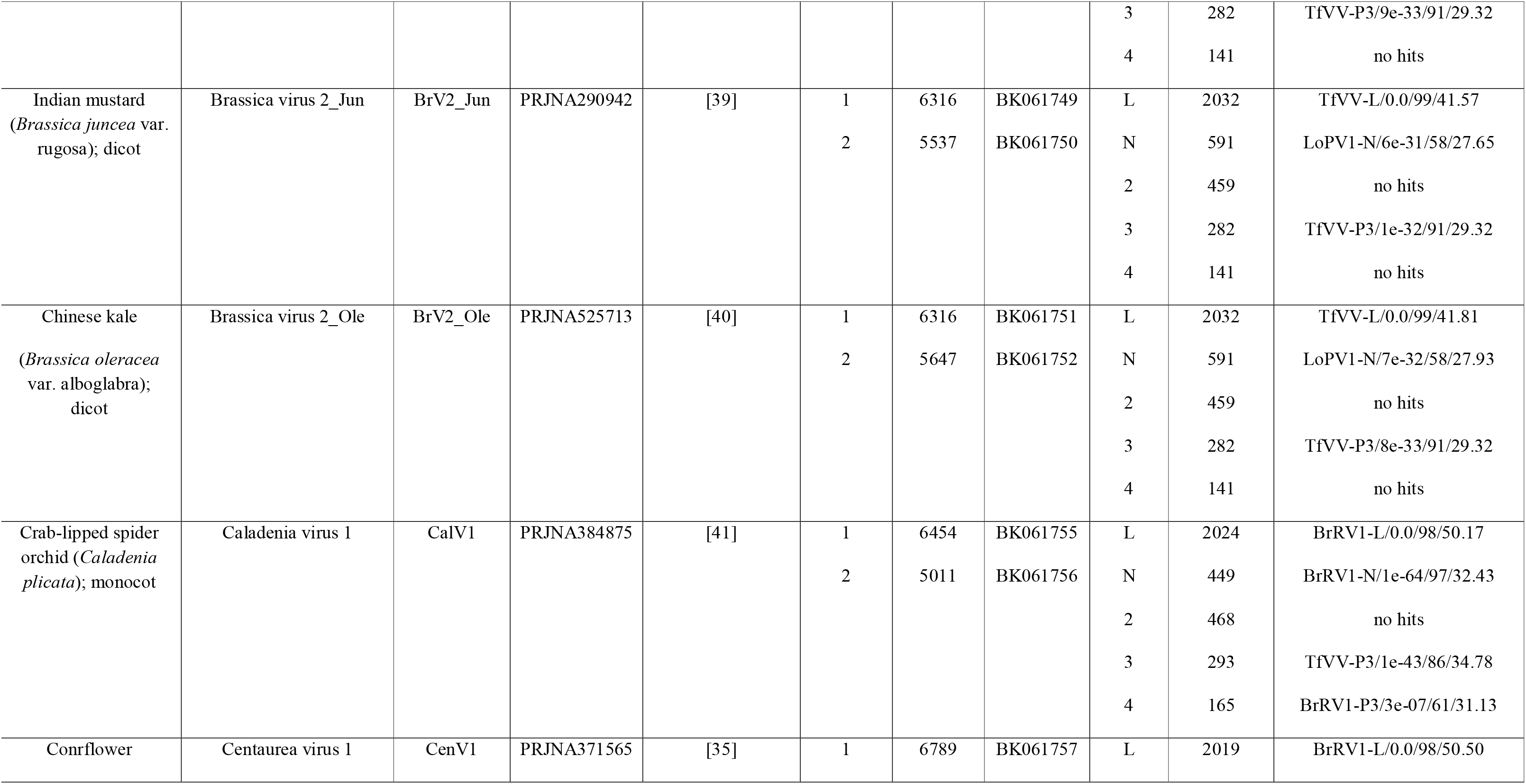
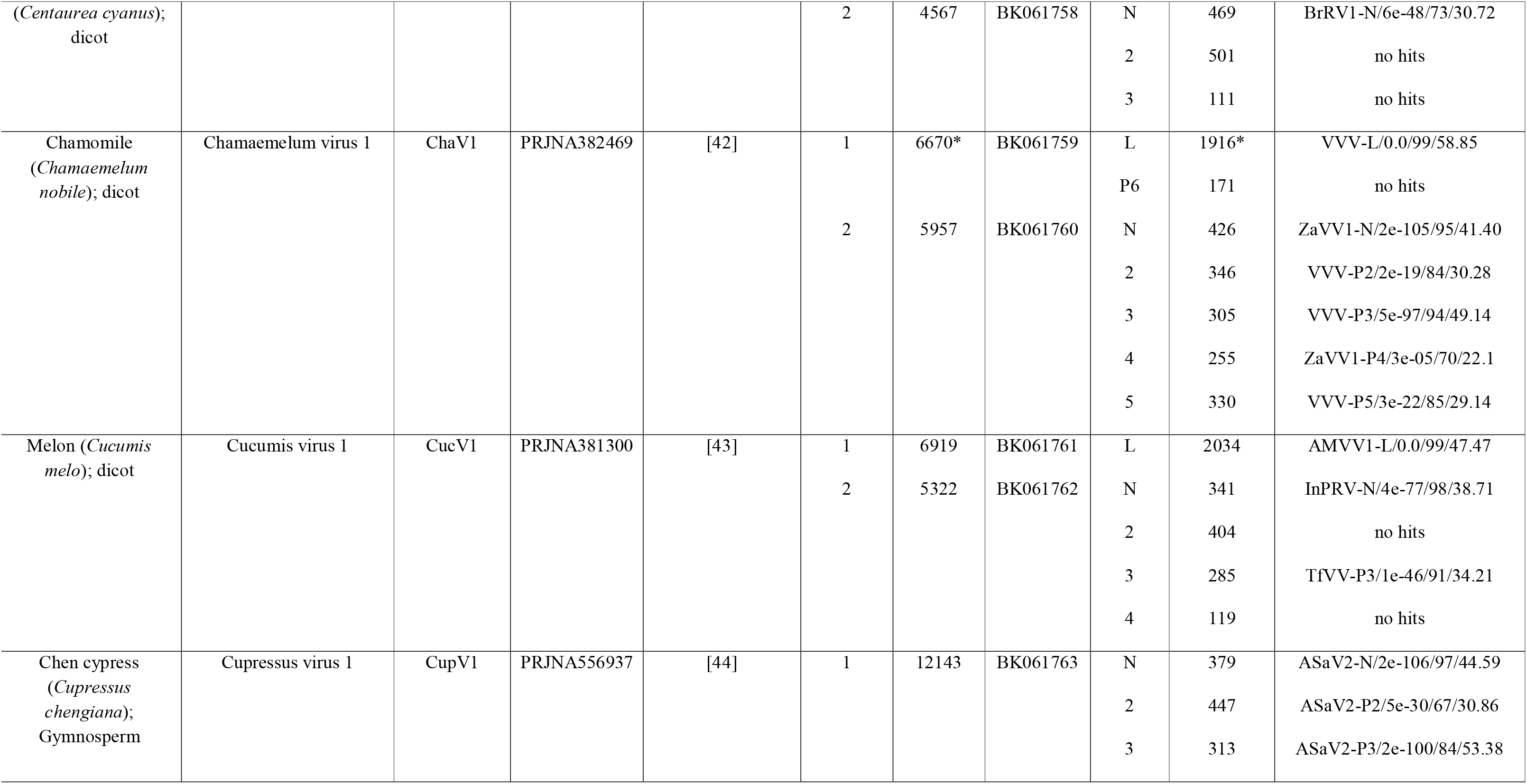
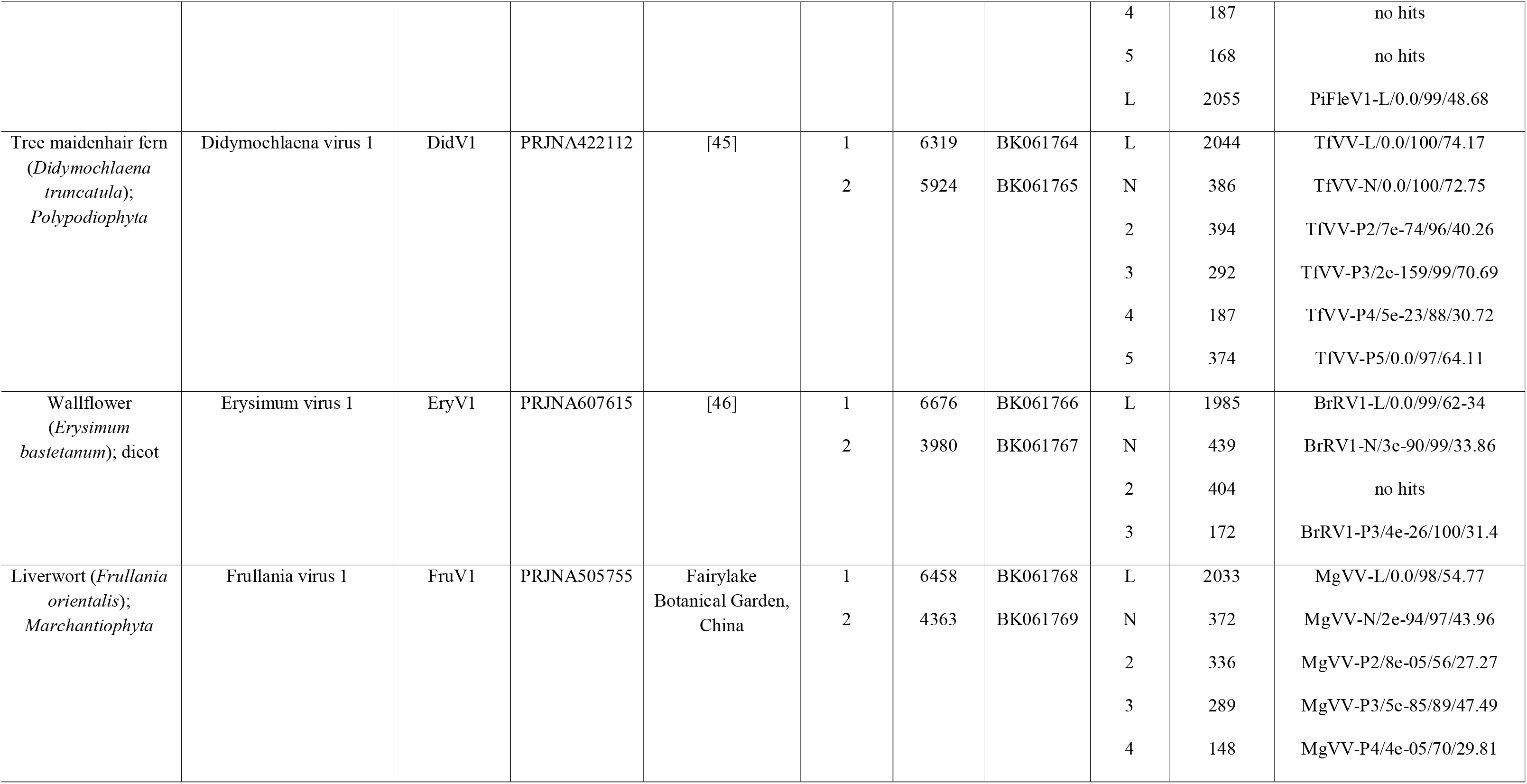
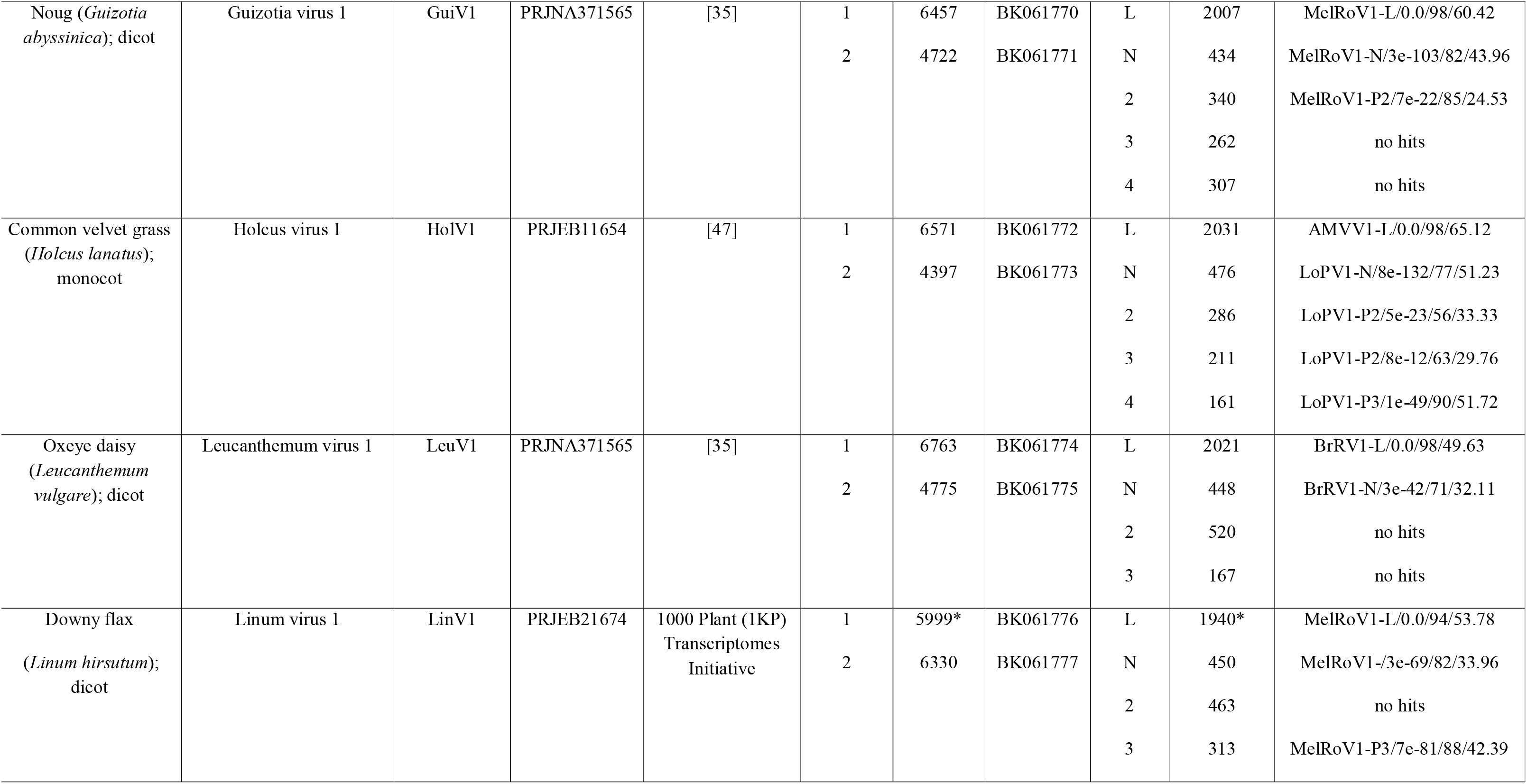
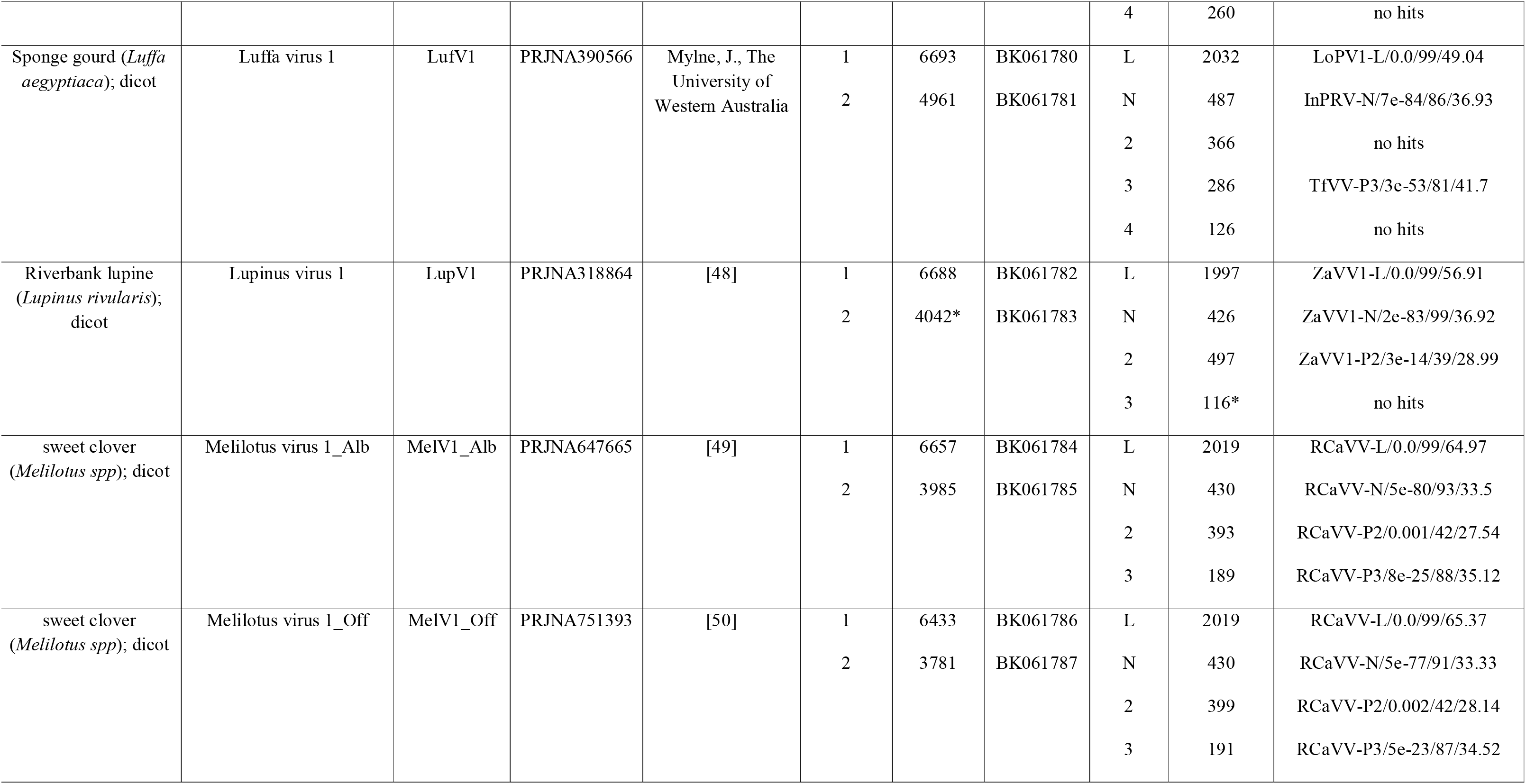
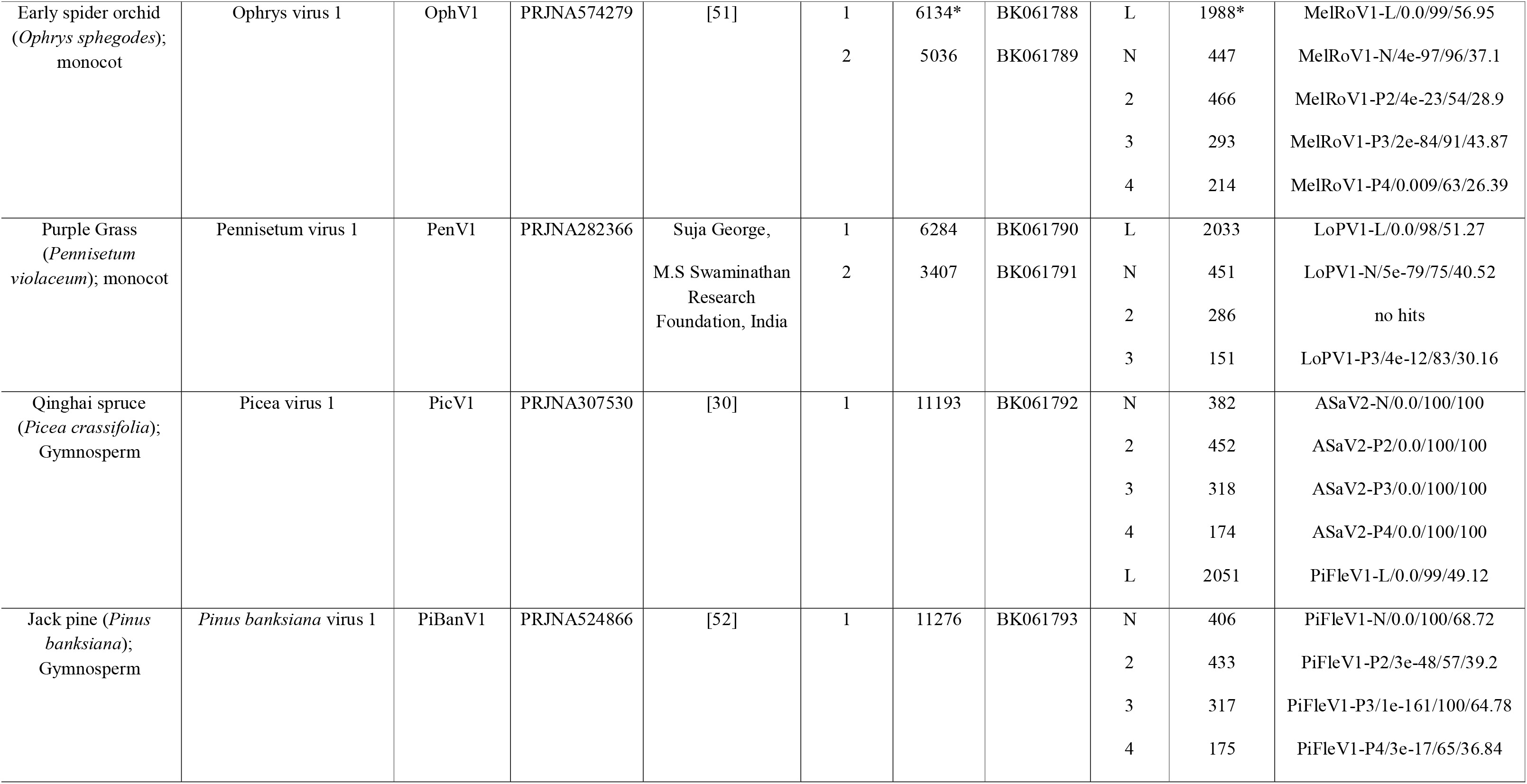
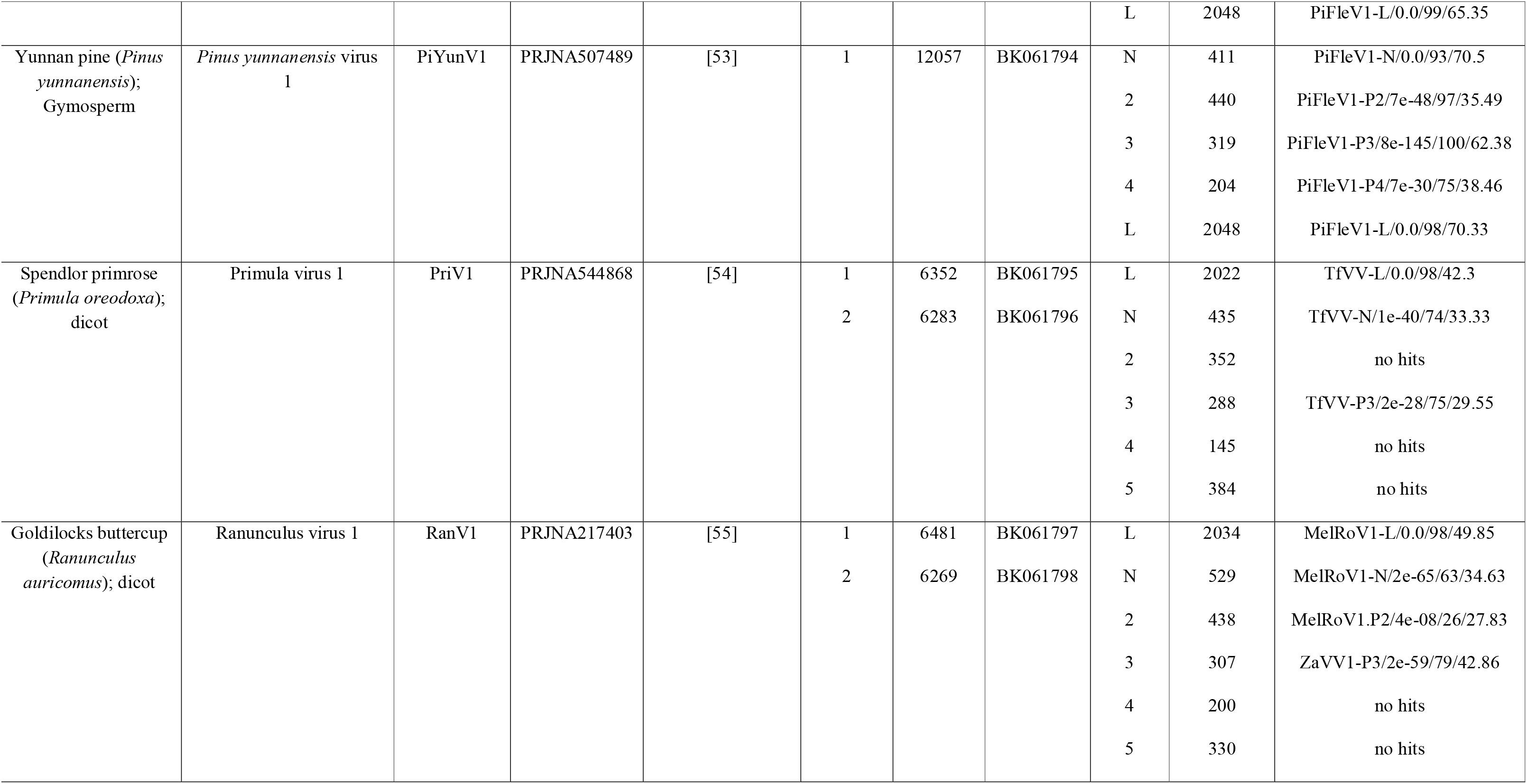
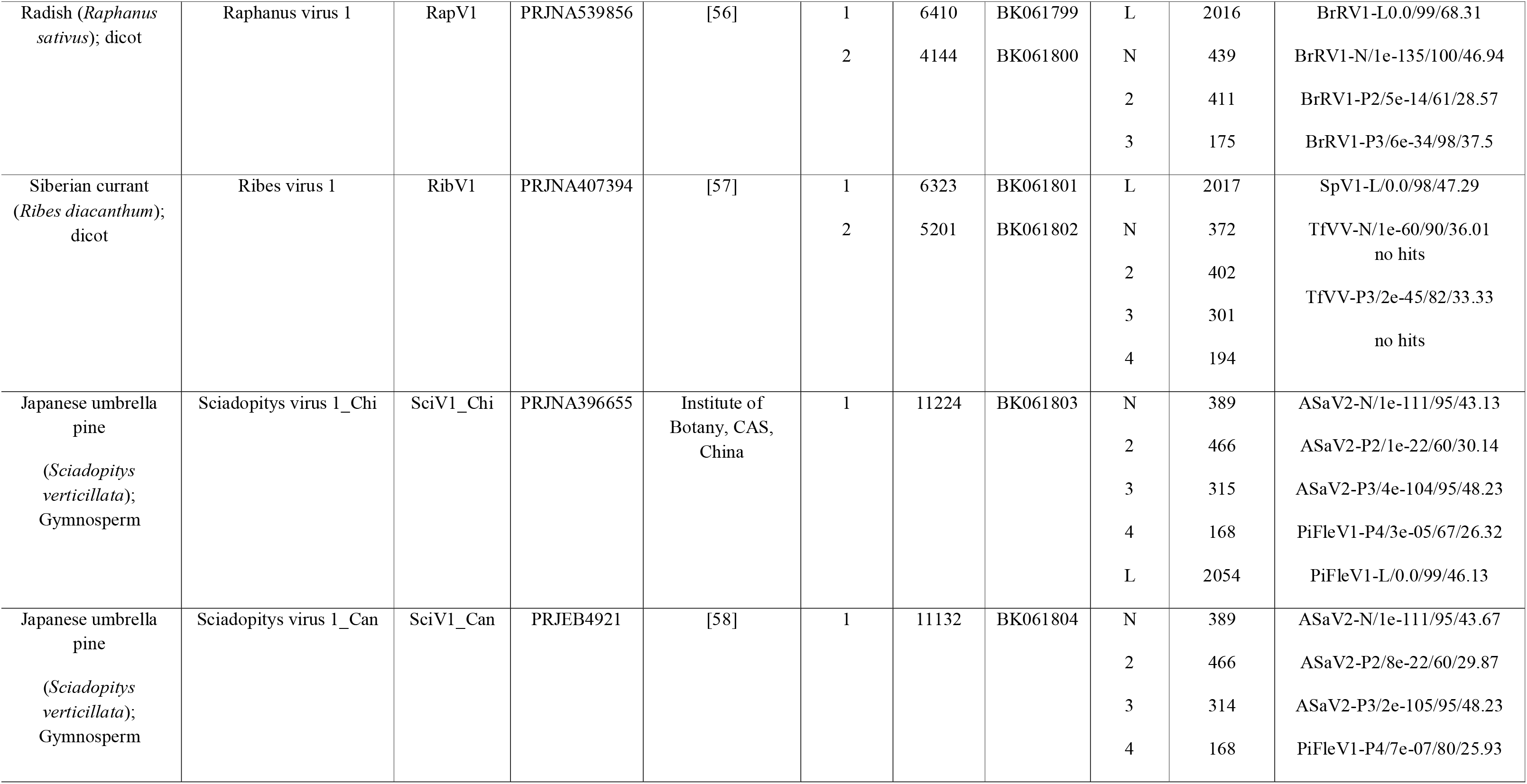
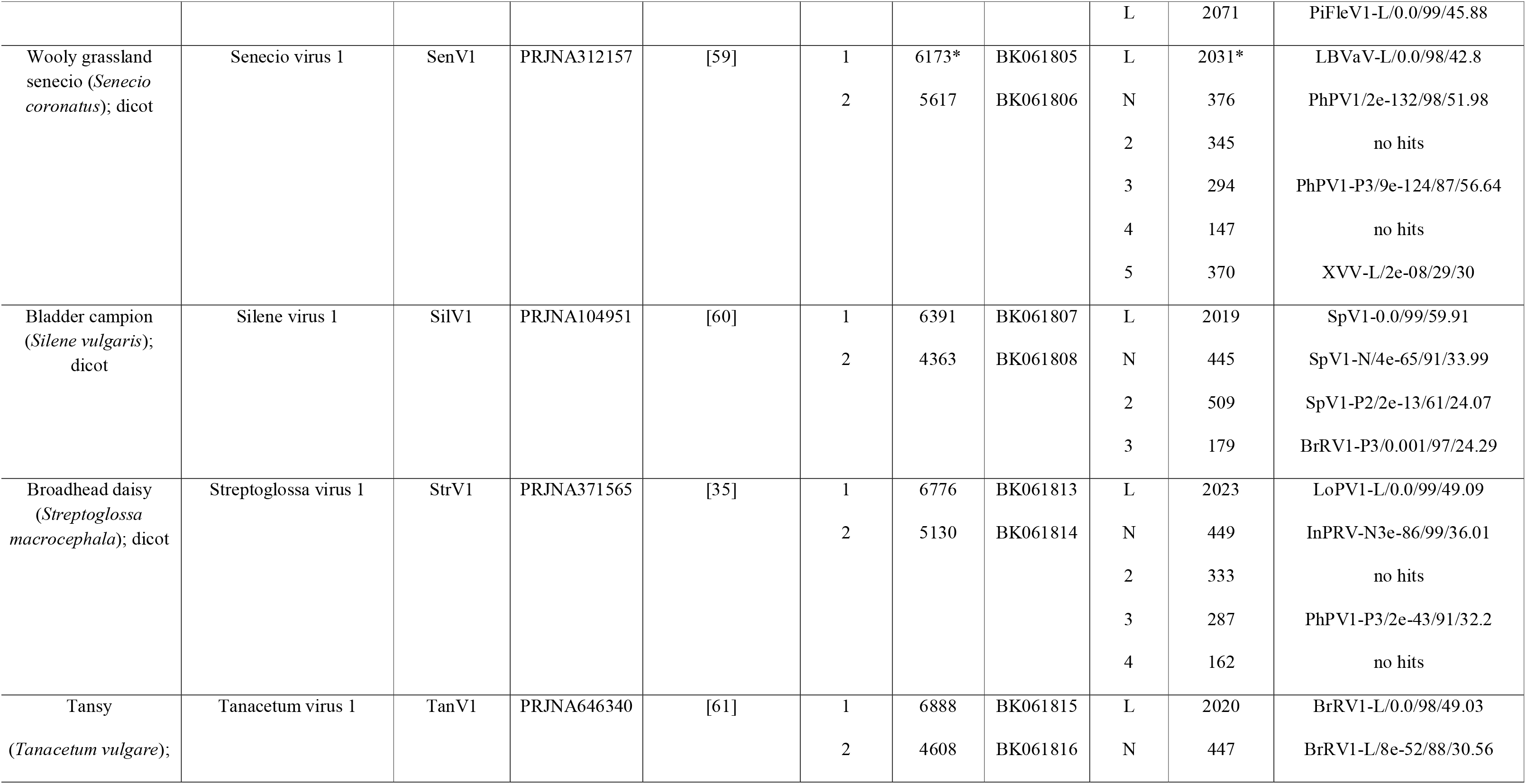
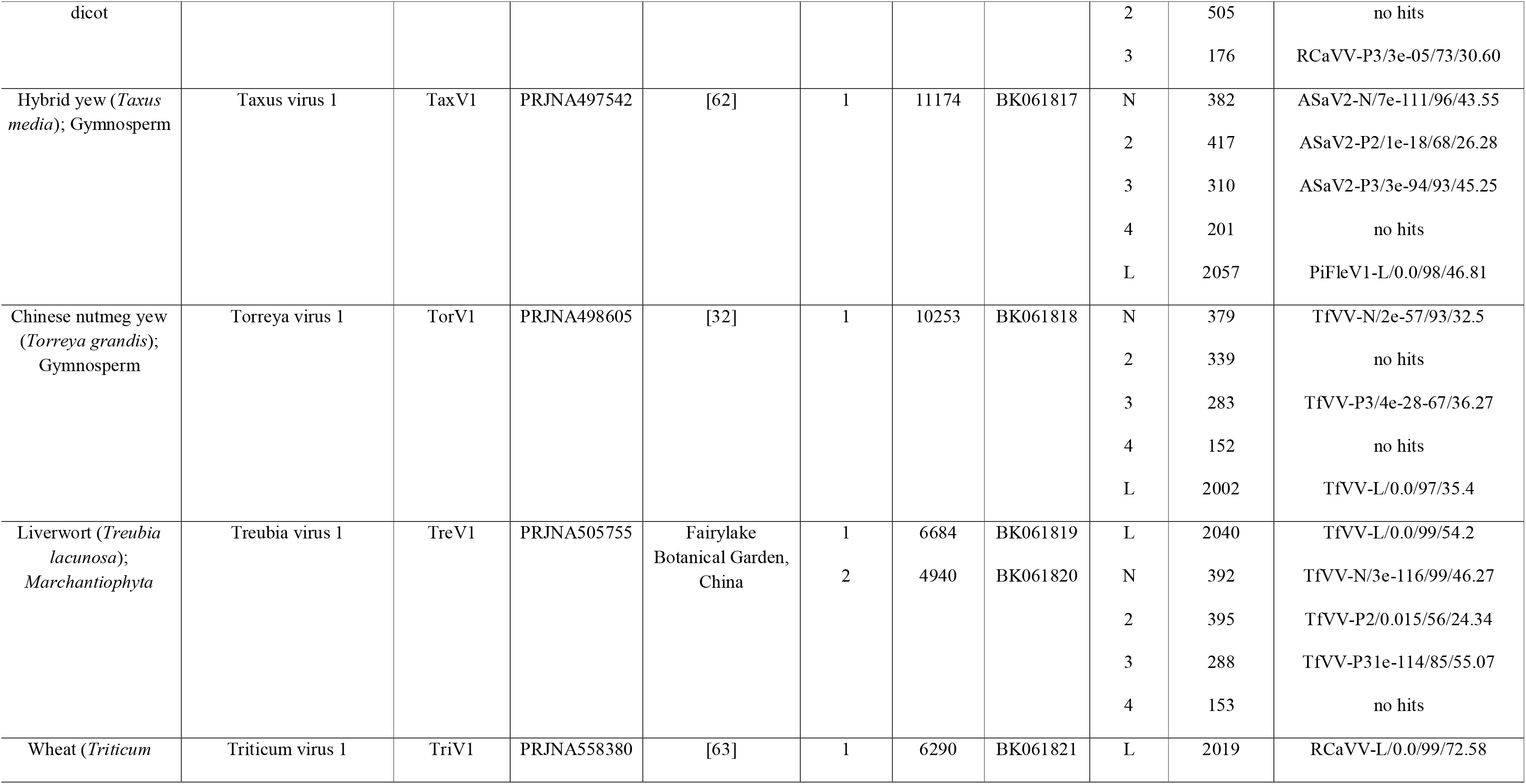
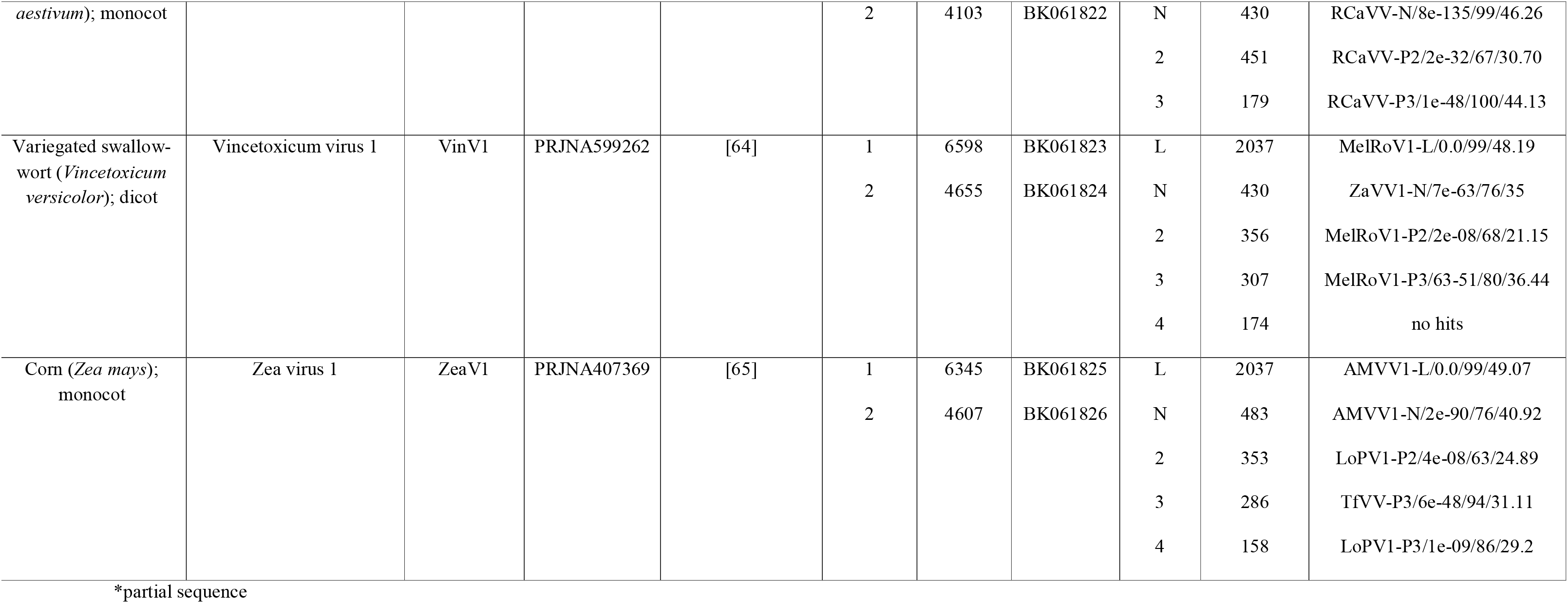
Summary of novel varicosaviruses identified from plant RNA-seq data available in the NCBI database. Acronyms of best hits are listed in Supp. Table 1

### 2.3 Bioinformatics tools and analyses

#### 2.3.1 Sequence analyses

ORFs were predicted with ORFfinder (minimal ORF length 150 nt, genetic code 1, https://www.ncbi.nlm.nih.gov/orffinder/), functional domains and architecture of translated gene products were determined using InterPro (https://www.ebi.ac.uk/interpro/search/sequence-search) and the NCBI Conserved domain database - CDD v3.19 (https://www.ncbi.nlm.nih.gov/Structure/cdd/wrpsb.cgi). Further, HHPred and HHBlits as implemented in https://toolkit.tuebingen.mpg.de/#/tools/ were used to complement annotation of divergent predicted proteins by hidden Markov models. Transmembrane domains were predicted using the TMHMM version 2.0 tool (http://www.cbs.dtu.dk/services/TMHMM/).

#### 2.3.2 Pairwise sequence identity

Percentage amino acid (aa) sequence identities of the L protein of those varicosaviruses identified in this study, as well as those available in the NCBI database were calculated using SDTv1.2 [23]. Virus names, abbreviations and NCBI accession numbers of varicosaviruses already reported are shown in Supp. Table 1.

#### 2.3.3 Phylogenetic analysis

Phylogenetic analysis based on the predicted polymerase protein of all available varicosaviruses was done using MAFFT 7.505 https://mafft.cbrc.jp/alignment/software with multiple aa sequence alignments using FFT-NS-i as the best-fit model. The aligned aa sequences were used as input to generate phylogenetic trees by the maximum-likelihood method (best-fit model = E-INS-i) with the FastTree 2.1.11 tool available at http://www.microbesonline.org/fasttree/. Local support values were calculated with the Shimodaira-Hasegawa test (SH) and 1,000 trees were resampled. L proteins of four selected cytorhabdoviruses were used as outgroup. To explore the potential phylogenetic co-divergence of varicosaviruses with their associated host plants, plant host cladograms were generated in phyloT v.2 (https://phylot.biobyte.de/), based on NCBI Taxonomy. Host associations were based on all varicosaviruses shown in Fig. 3. Connections were manually inferred between viral and plant phylogram and cladogram and visually inspected.

**Figure 1.**
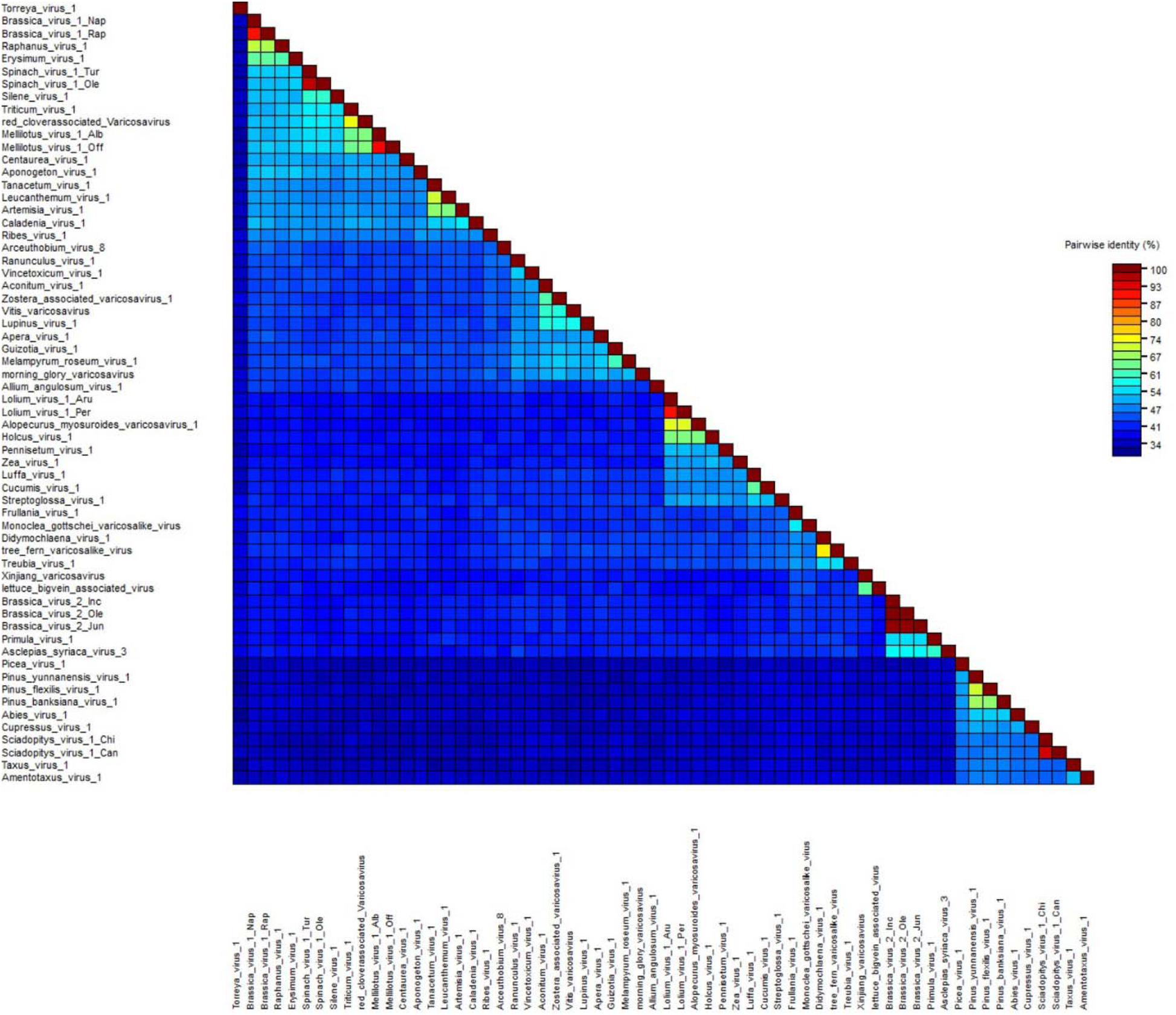
Pairwise identity matrix of the amino acid sequences of varicosavirus complete L gene open reading frame generated using SDT v1.2 software [23]. GenBank accession numbers are listed in Supplementary Table 1 and Table 1.

**Figure 2.**
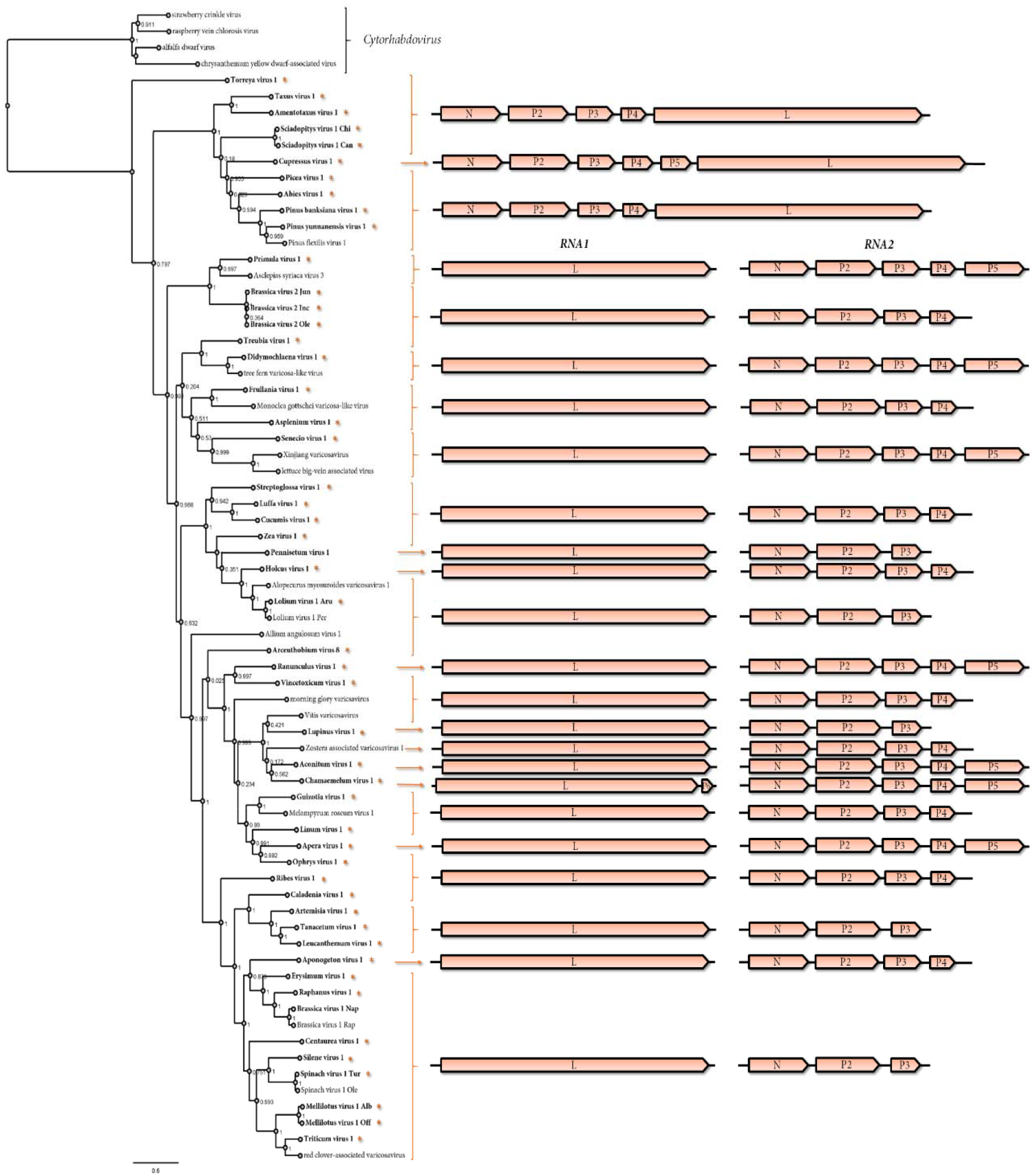
Left: Maximum-likelihood phylogenetic tree based on amino acid sequence alignments of the complete L gene of all varicosaviruses reported so far and in this study. The scale bar indicates the number of substitutions per site. Node labels indicate Fast Tree support values. Four cytorhabdoviruses were used as outgroup. Right: genomic organization of the varicosavirus sequences used in the phylogeny. An Asterisk and bold font indicate those viruses identified in this study. Accession numbers of all viruses are listed in Supplementary Table 1 and Table 1.

**Figure 3.**
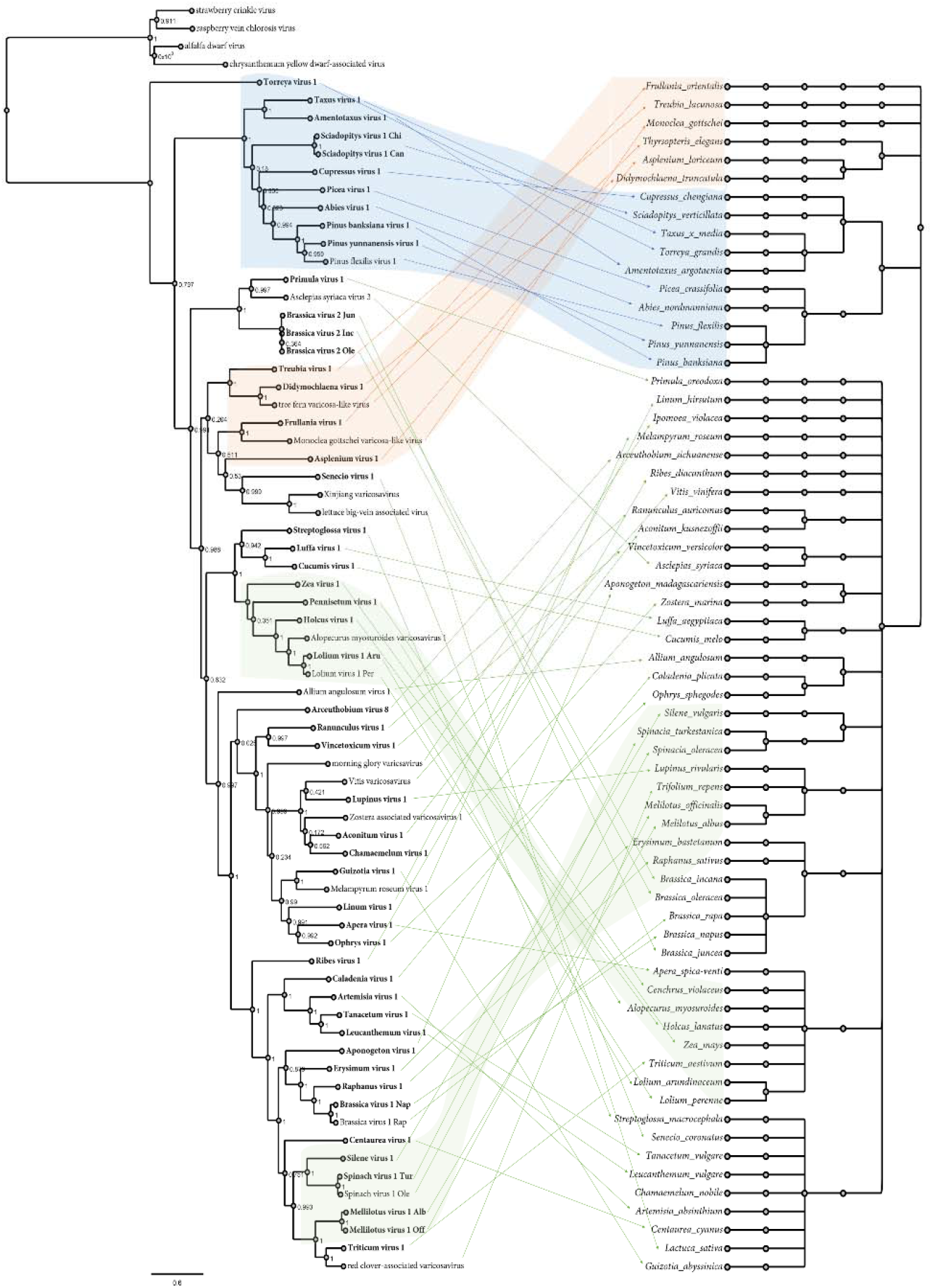
Tanglegram showing the phylogenetic relationships of varicosaviruses (left), which are linked with associated plant host(s) shown on the right. Links of well-supported clades of viruses to taxonomically related plant species are indicated in blue, orange, and green. Maximum likelihood phylogenetic tree of rhabdoviruses was constructed based on the conserved amino acid sequence of the complete L protein. Plant host cladograms were generated in phyloT v.2 based on NCBI Taxonomy. Internal nodes represent taxonomic structure of the NCBI taxonomy database including species, genus, family, order, subclass and sub-kingdom. Viruses identified in the present study are shown in bold font. The scale bar indicates the number of substitutions per site.

## 3. Results and Discussion

Most varicosaviruses likely do not induce easily discernable disease symptoms. Since their presence is not expected in sequencing libraries of apparently “healthy” vegetables, they are ideal candidates to be identified through mining of publicly available metatranscriptomic data. Accordingly, very recently, 12 novel proposed varicosaviruses were discovered when publicly available transcriptome datasets were mined [6, 9, 17–18]. Therefore, to unlock the hidden diversity of varicosaviruses, we extensively searched for these viruses in already available plant transcriptome data. This bioinformatics research resulted in the identification of 45 novel varicosaviruses, including the corrected full-length coding genome segments of the previously reported Arceuthobium sichuanense-associated virus 2 (ASaV2) [18] which had apparently been reconstructed from genome segments of two different varicosaviruses. We also identified three novel strains of three recently discovered varicosaviruses, confirming and strengthening results previously reported by Bejerman et al. [9]. This significant number of newly discovered varicosaviruses represents a 3.5-fold increase in the known varicosaviruses (Suppl. Fig. 1), which clearly highlights the importance of data-driven virus discovery to illuminate the landscape of largely overlooked taxonomic groups, such as the varicosaviruses.

More in detail, identification, assembly, and curation of raw SRA reads in this study resulted in 39 viral genome sequences with full-length coding regions and six with nearly complete coding regions. These viruses were associated with 45 plant host species (Table 1). Most of the tentative plant hosts of the novel varicosaviruses are herbaceous dicots (24/45), nine are herbaceous monocots, eight are gymnosperms and four are liverworts and ferns (Table 1).

The genomes of 37 viruses identified in this study were bisegmented, where RNA 1 of 36 of them encodes only the L protein, while RNA 1 of Chamaemelum virus 1 (ChaV1) has an additional ORF 5’ to the L gene, supported by the identification of the conserved intergenic sequence (see below), encoding a 171 aa putative protein (Table1, Fig. 2), which appears to be the first varicosavirus reported so far with an ORF in this position. RNA 2 segments of these 37 viruses have three to five genes in the order 3’-N-PX-5’. Twelve of them have three genes, while 17 have four genes and eight contained five genes (Table1, Fig. 2). Of the previously reported varicosaviruses six have three genes, four have four genes and four have five genes; therefore RNA 2 has a flexible genomic architecture and apparently the most frequent genomic organization in RNA 2 of varicosaviruses includes four genes (21 members) or three genes (18 members).

The consensus gene junction sequences of the bisegmented varicosaviruses was determined to be 3’ AU(N)_5_UUUUUGCUCU 5’ (Table 2), while the gene junction sequences of all, but one, unsegmented varicosavirus differed slightly in the 3’end being GU(N)_5_ instead of AU(N)_5_ (Table 2). Strikingly, the consensus gene junction of the unsegmented Torreya virus 1 (TorV1) was similar to that of bisegmented varicosaviruses. The potential implication of this difference in the gene junctions needs to be explored, since it could be linked to the basal evolutionary grouping of TorV1 (see below).

**Table 2.** Consensus varicosavirus gene junction sequences

| **Virus\*** | **3'end mRNA** | **intergenic spacer** | **5'end mRNA** |
| --- | --- | --- | --- |
| AbiV1 | CU (N) <sub>5</sub>UUUUU | G | CUCU |
| ArcV8 | AU (N) <sub>5</sub>UUUUU | G | CUCU |
| AcoV1 | AU (N) <sub>5</sub>UUUUU | G | CUCU |
| AmeV1 | CU (N) <sub>5</sub>UUUUU | G | CUCU |
| ApeV1 | AU (N) <sub>5</sub>UUUUU | G | CUCU |
| ApoV1 | AU (N) <sub>5</sub>UUUUU | G | CUCU |
| ArtV1 | AU (N) <sub>5</sub>UUUUU | G | CUCU |
| AscSyV3 | AU (N) <sub>5</sub>UUUUU | G | CUCU |
| AspV1 | AU (N) <sub>5</sub>UUUUU | G | CUCU |
| BrV2 | AU (N) <sub>5</sub>UUUUU | G | CUCU |
| CalV1 | AU (N) <sub>5</sub>UUUUU | G | CUCU |
| CenV1 | AU (N) <sub>5</sub>UUUUU | G | CUCU |
| ChaV1 | AU (N) <sub>5</sub>UUUUU | G | CUCU |
| CucV1 | AU (N) <sub>5</sub>UUUUU | G | CUCU |
| CupV1 | CU (N) <sub>5</sub>UUUUU | G | CUCU |
| DidV1 | AU (N) <sub>5</sub>UUUUU | G | CUCU |
| EryV1 | AU (N) <sub>5</sub>UUUUU | G | CUCU |
| FruV1 | AU (N) <sub>5</sub>UUUUU | G | CUCU |
| GuiV1 | AU (N) <sub>5</sub>UUUUU | G | CUCU |
| HolV1 | AU (N) <sub>5</sub>UUUUU | G | CUCU |
| LeuV1 | AU (N) <sub>5</sub>UUUUU | G | CUCU |
| LinV1 | AU (N) <sub>5</sub>UUUUU | G | CUCU |
| LufV1 | AU (N) <sub>5</sub>UUUUU | G | CUCU |
| LupV1 | AU (N) <sub>5</sub>UUUUU | G | CUCU |
| MelV1 | AU (N) <sub>5</sub>UUUUU | G | CUCU |
| OphV1 | AU (N) <sub>5</sub>UUUUU | G | CUCU |
| PenV1 | AU (N) <sub>5</sub>UUUUU | G | CUCU |
| PicV1 | CU (N) <sub>5</sub>UUUUU | G | CUCU |
| PiBanV1 | CU (N) <sub>5</sub>UUUUU | G | CUCU |
| PiYunV1 | CU (N) <sub>5</sub>UUUUU | G | CUCU |
| Priv1 | AU (N) <sub>5</sub>UUUUU | G | CUCU |
| RanV1 | AU (N) <sub>5</sub>UUUUU | G | CUCU |
| RapV1 | AU (N) <sub>5</sub>UUUUU | G | CUCU |
| RibV1 | AU (N) <sub>5</sub>UUUUU | G | CUCU |
| SciV1 | CU (N) <sub>5</sub>UUUUU | G | CUCU |
| SenV1 | AU (N) <sub>5</sub>UUUUU | G | CUCU |
| SilV1 | AU (N) <sub>5</sub>UUUUU | G | CUCU |
| StrV1 | AU (N) <sub>5</sub>UUUUU | G | CUCU |
| TanV1 | AU (N) <sub>5</sub>UUUUU | G | CUCU |
| TaxV1 | CU (N) <sub>5</sub>UUUUU | G | CUCU |
| TorV1 | AU (N) <sub>5</sub>UUUUU | G | CUCU |
| TreV1 | AU (N) <sub>5</sub>UUUUU | G | CUCU |
| TriV1 | AU (N) <sub>5</sub>UUUUU | G | CUCU |
| VinV1 | AU (N) <sub>5</sub>UUUUU | G | CUCU |
| ZeaV1 | AU (N) <sub>5</sub>UUUUU | G | CUCU |
| AAnV1 | AU (N) <sub>5</sub>UUUUU | G | CUCU |
| AMVV1 | AU (N) <sub>5</sub>UUUUU | G | CUCU |
| BrV1 | AU (N) <sub>5</sub>UUUUU | G | CUCA |
| LBVaV | AU (N) <sub>5</sub>UUUUU | G | CUCU |
| LoV1 | AU (N) <sub>5</sub>UUUUU | G | CUCU |
| MelRoV1 | AU (N) <sub>5</sub>UUUUU | G | CUCU |
| MGVV | AU (N) <sub>5</sub>UUUUU | G | CUCU |
| MgVV | AU (N) <sub>5</sub>UUUUU | G | CUCU |
| PhPiV1 | AU (N) <sub>5</sub>UUUUU | G | CUCU |
| PiFleV1 | GU (N) <sub>5</sub>UUUUU | G | CUCU |
| RCaVV | AU (N) <sub>5</sub>UUUUU | G | CUCU |
| SpV1 | AU (N) <sub>5</sub>UUUUU | G | CUCU |
| TfVV | AU (N) <sub>5</sub>UUUUU | G | CUCU |
| VVV | AU (N) <sub>5</sub>UUUUU | G | CUCU |
| XVV | AU (N) <sub>5</sub>UUUUU | G | CUCU |
| ZaVV1 | AU (N) <sub>5</sub>UUUUU | G | CUCU |

There is a great dearth regarding data on the potential functions of putative proteins, other than N and L, encoded by varicosaviruses, and intriguingly, no conserved domains were identified in these proteins. We grasped some shared identities mostly for the cognate P3, but also for several P2 proteins (Table 1), but for most encoded proteins BlastP results were orphans, no known signals or domains present, and not a single clue towards their putative (conserved?) function. Thus, further studies should be focused on the functional characterization of these proteins to gain essential knowledge regarding the elusive proteome of varicosaviruses beyond N and L proteins.

The pairwise aa sequence identities between the L proteins of all reported varicosaviruses, including those identified in this study, showed a great diversity and an overall low identity between the different varicosaviruses (Fig.1, Supp. Table S2). Relatively low sequence identity is a common feature among rhabdovirus taxa, characterized by a high level of diversity in both genome sequence and organization [10]. In addition, the overall low sequence identity among the novel viruses detected here and with the previously described varicosaviruses suggests that despite the many viruses identified in this study, there is likely still a significant amount of virus “dark matter” of yet to be discovered varicosaviruses.

When we analyzed the diversity between strains of viruses which are likely members of the same species, we found that proteins encoded by Brassica virus 2, Spinach virus 1 and Sciadopitys virus 1 strains were very similar. On the other hand, proteins encoded by Brassica virus 1, Lolium virus 1 and Melilotus virus 1 strains were quite diverse, but nevertheless showing aa identities for N and L proteins exceeding 80%. Thus, we tentatively propose an aa sequence identity of 80% across the L gene as threshold for species demarcation in the *Varicosavirus* genus, a taxonomic criterion which had previously not been fully defined [10]. This threshold is strongly supported by the comparison of the L protein aa sequence of 60 viruses (Fig.1, Suppl. Table 2). Based on this criterion all 39 novel viruses with their complete coding region assembled in this study should be considered as belonging to novel *Varicosavirus* species, which would increase the number of members of the genus by more than an order of magnitude.

Bejerman et al. [9] tentatively reported the first unsegmented varicosavirus, Pinus flexilis virus 1 (PiFleV1) which was associated with the *gymnosperm Pinus flexilis*. In this study we complemented that result by the discovery of eight additional unsegmented varicosaviruses, which were exclusively associated with gymnosperms (Table 1), some of them linked to the same genus *Pinus* and presenting a significant co-evolution of viruses and hosts. These results robustly support a clade of gymnosperm-associated varicosaviruses with a distinct genome architecture, requiring the rewriting of a previously proposed key feature and fundamental marker of varicosaviruses; their genomic bisegmented nature. It is tempting to speculate that the unsegmented genomic architecture may be linked to the adaptation to gymnosperm hosts and a shared ancient evolutionary history of these viruses and hosts.

Interestingly, in BlastP analyses of N, P2 and P3 of those gymnosperm-associated viruses, most of them had as best hit to the cognate proteins encoded by the putative bisegmented ASaV2 (Table 1) a virus apparently hosted by a parasitic plant of spruce (*Picea, Pinacea*). Furthermore, unexpectedly the best hit of the putative P5 protein encoded on ASaV2 RNA2 was a fragment of the PiFleV1 L protein, while the deduced L protein on ASaV2 RN1 was not a best hit with PiFleV1, but instead with the non-gymnosperm linked MelRoV1 hosted by the Orobanchaceae parasitic plant*Melampyrum roseum*. Thus, we suspected that ASaV2 was potentially miss-assembled from fragments belonging to two different viruses. Consequently, we re-analyzed the original SRA data used by Sidhartan et al. [18] and were able to assemble two distinct varicosavirus genomes, one bisegmented genome presumably linked to the parasitic plant and one unsegmented genome most probably linked to spruce, which would support our hypothesis. We believe that there are several reasons that led to the original ASaV2 description: *i*) the atypical and unexpected existence at the time of an unsegmented varicosavirus. *ii*) the presence of two varicosaviruses in the very same sequencing library, which may be the first tentative evidence in the literature of coinfection of two varicosaviruses. *iii*) the fact that the sequence reads corresponding to the L gene region of the unsegmented varicosavirus where low which may have affected the assembling pipelines used by the authors. All in all, independently verifying unexpected re-analysed SRA data may lead to a clearer understanding of the genomic structure of mined RNA virus genomes.

The phylogenetic analysis based on deduced L protein aa sequences placed all unsegmented varicosaviruses, except TorV1, into a distinct clade. Interestingly TorV1 was placed in a clade that was basal to the other unsegmented varicosaviruses (Fig.2). This distinct phylogenetic branching and clustering of the unsegmented viruses suggests that they share a unique evolutionary history among varicosaviruses. Bisegmented varicosaviruses did not cluster according to their genomic organization nor with the plant species associated with each virus (Fig. 2). For example, brassica virus 1 and brassica virus 2 belong to distinct clades, while two viruses associated with orchids (Ophius virus 1 and Caladenia virus 1) were also placed in different clusters, as well as monocot-associated viruses did not all group together. On the other hand, all varicosaviruses associated with ferns and liverworts belonged to the same cluster, also shared with previously reported varicosaviruses from these plant types, while most of the grass-associated varicosaviruses also clustered together (Fig. 2).

We generated a tanglegram to compare the virus phylogram and plant host cladogram to further explore virus-host relationships (Fig. 3). This analysis showed that viruses of some clades clearly co-diverged with their hosts, including the gymnosperm-associated virus clade, the SpV1 and Silene virus 1 clade, the grass-associated virus clade, and the clade of fern and liverworts viruses, suggesting a shared host-virus evolution in those clades (Fig. 3). However, the tanglegram topology also indicated that for most of the varicosaviruses, there is no apparent concordant evolutionary history with their plant hosts, similar to what was previously reported for invertebrate and vertebrates rhabdoviruses [24].

Several lines of evidence suggest that varicosaviruses may be vertically transmitted; i) a close host-virus co-evolution in some clades may reflect species isolation and a lack of horizontal transmission, ii) some viruses detected in this study were identified from seed transcriptomics databases, iii) an emerging characteristic of persistent, chronic infections of several plant viruses which are likely vertically transmitted are latent/asymptomatic infections, a characteristic which appears to be shared with varicosaviruses. Thus, further studies should be carried out to elucidate the transmission mode of varicosaviruses beyond the fungal-transmitted LBVaV [11]. It is worth mentioning that even with the availability of thousands of RNAseq libraries of fungi or arthropods, we failed to detect any evidence of varicosaviruses in those organisms, which could suggest that vectors of varicosaviruses are rare or non-existent.

Before the era of data-driven virus discovery, few viruses had been identified in gymnosperms [25–27]. However, when data mining was applied to publicly available transcriptomes, many novel viruses were identified in this large group of higher plants, highlighting the rich and diverse gymnosperms virosphere, which still is largely unexplored. A distinct clade of gymnosperm-associated viruses was also recently identified within amalgaviruses [28]; while we recently described two distinct caulimovirids and geminivirids linked to the gnetophyte *Welwitschia mirabilis* [29]. Eight unsegmented varicosaviruses associated with gymnosperms were identified in this study, and another was discovered by Bejerman et al. [9]. Taken together, all these viruses recently discovered in gymnosperms strongly suggests that they may have evolutionary trajectories that are distinct from those infecting angiosperms. Thus, it is likely that further exploration of additional gymnosperm datasets or new transcriptome studies of other gymnosperms will yield plenty of novel viruses with unique features highlighting the close evolution with their hosts. The clear association between gymnosperm-associated viruses and their hosts likely indicates a close coevolution, which suggest an early adaptation of this group of viruses to infect gymnosperms. This hypothesis is also supported by the distinct genomic architecture and divergent evolutionary history among varicosaviruses as shown in the phylogenetic tree, characterized by long branches and distinctive clustering. Taken together, the gymnosperm-associated varicosaviruses could be taxonomically classified in a novel genus within the family *Rhabdoviridae*, subfamily *Betarhabdovirinae* for which we suggest the name “*Gymnorhavirus*”.

In summary, this study highlights the importance of the analysis of SRA public data as a valuable tool not only to accelerate the discovery of novel viruses but, also, to gain insight into their evolution and to refine virus taxonomy. Using this approach, we looked for hidden varicosa-like virus sequences to unlock the veiled diversity of a largely neglected plant rhabdovirus genus, the varicosaviruses. Our findings, including an about 3.5-fold expansion of the current genomic diversity within the genus, resulted in the most complete phylogeny of varicosaviruses to date and shed new light on the genomic architecture, phylogenetic relationships and evolutionary landscape of this unique group of plant rhabdoviruses. Future studies should assess many intriguing aspects on the biology and ecology of these viruses such as potential symptoms, vertical transmission, and putative vectors.

## Acknowledgments

We would like to express sincere gratitude to the generators of the underlying data used for this work, which are cited in Table 1. By following open-access practices and supporting accessible raw sequence data in public repositories available to the research community, they promoted the generation of new knowledge and ideas.

## Author Contributions

Conceptualization, N.B, R.D and H.D; data analysis, N.B and H.D; writing—original draft preparation, N.B; writing—review and editing, N.B, R.D and H.D. All authors have read and agreed to the published version of the manuscript.

## Institutional Review Board Statement

Not applicable for studies not involving humans or animals.

## Informed Consent Statement

Not applicable for studies not involving humans.

## Data availability statement

Nucleotide sequence data reported are available in the Third Party Annotation Section of the DDBJ/ENA/GenBank databases under the accession numbers TPA: BK061731-BK061826. These sequences are available as Supplementary Materials.

## References

1. Koonin, E.V., Krupovic, M., Agol, V.I. (2021). The Baltimore Classification of Viruses 50 Years Later: How Does It Stand in the Light of Virus Evolution? Microbiol. Mol. Biol. Rev. 85:e0005321.

2. Koonin, E.V., Dolja, V.V., Krupovic, M., Varsani, A., Wolf, Y.I., Yutin, N. et al. (2020). Global Organization and Proposed Megataxonomy of the Virus World. Microbiol Mol. Biol. Rev. 84:1–33.

3. Geoghegan, J.L., Holmes, E.C. (2017a). Predicting virus emergence amid evolutionary noise. Open Biol. 7:170189.

4. Dolja, V.V., Krupovic, M., Koonin, E.V. (2020). Deep roots and splendid boughs of the global plant virome. Annu. Rev. Phytopathol. 58:23–53.

5. Edgar, R.C., Taylor, J., Lin, V., Altman, T., Barbera, P., et al. (2022) Petabase-scale sequence alignment catalyses viral Discovery. Nature. 602:142–147.

6. Mifsud, J., Gallagher, R., Holmes, E., Geoghegan, J. (2022). Transcriptome mining expands knowledge of RNA viruses across the plant kingdom. J Virol. e0026022.

7. Lauber, C., Seitz, S. (2022). Opportunities and Challenges of Data-Driven Virus Discovery. Biomolecules 12:1073.

8. Dietzgen R. G., Bejerman N. E., Goodin M. M., Higgins C. M., Huot O. B., Kondo H., et al. (2020). Diversity and epidemiology of plant rhabdoviruses. Virus Res. 281:197942.

9. Bejerman, N., Dietzgen, R.G., Debat, H. (2021). Illuminating the Plant Rhabdovirus Landscape through Metatranscriptomics Data. Viruses 13:1304.

10. Walker, P.J., Freitas-Astúa, J., Bejerman, N., Blasdell, K.R., Breyta, R., et al. (2022). ICTV Virus Taxonomy Profile: Rhabdoviridae 2022. J. Gen. Virol. 103:001689.

11. Campbell, R. N. (1996). Fungal transmission of plant viruses. Annu. Rev. Phytopathol. 34:87–108.

12. Sasaya, T., Ishikawa, K., and Koganezawa, H. (2002). The nucleotide sequence of RNA1 of Lettuce big-vein virus, genus Varicosavirus, reveals its relation to nonsegmented negative-strand RNA viruses. Virology 297:289–297.

13. Sasaya, T., Kusaba, S., Ishikawa, K., Koganezawa, H., (2004). Nucleotide sequence of RNA2 of *Lettuce big-vein virus* and evidence for a possible transcription termination/initiation strategy similar to that of rhabdoviruses. J Gen Virol. 85:2709–17.

14. Verbeek, M., Dullemans, A.M., van Bekkum, P.J., van der Vlugt, R.A.A. (2013). Evidence for Lettuce big-vein associated virus as the causal agent of a syndrome of necrotic rings and spots in lettuce. Plant Pathol. 62:444–451.

15. Koloniuk, I., Fránová, J., Sarkisova, T., Přibylová, J., Lenz, O., et al. (2018). Identification and molecular characterization of a novel varicosa-like virus from red clover. Arch. Virol. 163:2213–2218.

16. Sabbadin, F., Glover, R., Stafford, R., Rozado-Aguirre, Z., Boonham, N., et al. (2017). Transcriptome sequencing identifies novel persistent viruses in herbicide resistant wild-grasses. Sci. Rep. 7:41987.

17. Shin, C., Choi, D., Hahn, Y. (2021). Identification of the genome sequence of Zostera associated varicosavirus 1, a novel negative-sense RNA virus, in the common eelgrass (Zostera marina) transcriptome. Acta Virol, 65:373–380.

18. Sidharthan, V.K., Chaturvedi, K.K., Baranwal, V.K. (2022a). Diverse RNA viruses in a parasitic owering plant (spruce dwarf mistletoe) revealed through RNA-seq data mining. J Gen Plant Pathol 88:138–44.

19. Chen, Y., Sadiq, S., Tian, J., Chen, X., Lin, X., et al. (2022). RNA viromes from terrestrial sites across China expand environmental viral diversity. Nat Microbiol. 7:1312–1323.

20. Nabeshima, T., Abe, J. (2021). High-throughput sequencing indicates novel Varicosavirus, Emaravirus and Deltapartitvirus infections in *Vitis coignetiae*. Viruses 13:827.

21. Zhao, F., Liu, H., Qiao, Q, Wang, Y., Zhang, D., et al. (2021). Complete genome sequence of a novel varicosavirus infecting tall morning glory (Ipomoea purpurea). Arch Virol. 166:3225–3228.

22. Leebens-Mack, J.H., Barker, M.S., Carpenter, E.J., et al. (2019). One thousand plant transcriptomes and the phylogenomics of green plants. Nature 574:679–685.

23. Muhire, B.M., Varsani, A., Martin, D.P. (2014). SDT: a virus classification tool based on pairwise sequence alignment and identity calculation. PLoS One 9:e108277.

24. Geoghegan, J. L., Duchêne, S., Holmes, E. C. (2017b) Comparative analysis estimates the relative frequencies of co-divergence and cross-species transmission within viral families. PLOS Pathog. 13:e1006215.

25. Alvarez-Quinto, R., Lockhart, B., Fetzer, J., Olszewski, N. (2020). Genomic characterization of cycad leaf necrosis virus, the first badnavirus identified in a gymnosperm. Arch Virol. 165:1671–1673.

26. Koh, S.H., Li, H., Admiraal, R., Jones, M.G.K., Wylie, S.J. (2015). Catharanthus mosaic virus: A potyvirus from a gymnosperm, *Welwitschia mirabilis*. Virus Res. 203:41–46.

27. Rastrojo, A., Núñez, A., Moreno, D.A., Alcamí, A. (2018) A new putative Caulimoviridae genus discovered through air metagenomics. Microbiol. Resour. Announc. 7:e00955–18.

28. Sidharthan, V.K., Rajeswari, V., Vanamala, G., Baranwal, V.K. (2022b). Revisiting the amalgaviral landscapes in plant transcriptomes expands the host range of plant amalgaviruses. https://doi.org/10.21203/rs.3.rs-2012542/v1.

29. Debat, H., Bejerman, N (2022). A glimpse into the DNA virome of the unique “living fossil” Welwitschia mirabilis. Gene. 843:146806.

30. Wang, Y., Li, X., Zhou, W., Li, T., Tian, C. (2016). De novo assembly and transcriptome characterization of spruce dwarf mistletoe Arceuthobium sichuanense uncovers gene expression profiling associated with plant development. BMC Genomics. 17:771.

31. Tang, M., Zhao, W., Xing, M., Zhao, J., Jiang, Z., et al. (2021). Resource allocation strategies among vegetative growth, sexual reproduction, asexual reproduction and defense during growing season of *Aconitum kusnezoffii* Reichb. Plant J. 105:957–977.

32. Yu, C., Zhan, X., Zhang, C., Xu, X., Huang, J., Feng, S., et al. (2021a). Comparative metabolomic analyses revealed the differential accumulation of taxoids, flavonoids and hormones among six Taxaceae trees. Sci. Hortic. 285:110196.

33. Babineau, M., Mahmood, K., Mathiassen, S.K., Kudsk, P., Kristensen, M. (2017). *De novo* transcriptome assembly analysis of weed *Apera spica-venti* from seven tissues and growth stages. BMC Genom. 18:128.

34. Rowarth, N., Curtis, B., Einfeldt, A., Archibald, J., Lacroix, C., Gunawardena, A. (2021). RNA-Seq analysis reveals potential regulators of programmed cell death and leaf remodelling in lace plant (Aponogeton madagascariensis). BMC Plant Biol. 21:375

35. Jayasena, A.S., Fisher, M.F., Panero, J.L., Secco, D., Bernath-Levin, K., et al. (2017). Stepwise evolution of a buried inhibitor peptide over 45 My. Mol. Biol. Evol. 34:1505–1516.

36. Weitemier, K., Straub, S., Fishbein, M., Bailey, C., Cronn, R., Liston, A. (2019). A draft genome and transcriptome of common milkweed *(Asclepias syriaca)* as resources for evolutionary, ecological, and molecular studies in milkweeds and Apocynaceae. PeerJ 7: e7649.

37. Shen, H.., Jin, D.., Shu, J.-P., Zhou, X.-L., Lei, M., et al. (2018). Large-scale phylogenomic analysis resolves a backbone phylogeny in ferns. Gigascience 7:1–11.

38. An, H., Qi, X., Gaynor, M. L., Hao, Y., Gebken, S. C., et al. (2019). Transcriptome and organellar sequencing highlights the complex origin and diversification of allotetraploid Brassica napus. Nat. Commun. 10:2878.

39. Bisht, D.S., Chamola, R., Nath, M., Bhat, S.R. (2015). Molecular mapping of fertility restorer gene of an alloplasmic CMS system in *Brassica juncea* containing Moricandia arvensis cytoplasm. Mol. Breed. 35: 14.

40. Wu, Q., Wang, J., Mao, S., Xu, H., Wu, Q., et al. (2019). Comparative transcriptome analyses of genes involving in sulforaphane metabolism at different treatment in Chinese kale using full-length transcriptome sequencing. BMC Genom. 20:377.

41. Xu, H., Bohman, B., Wong, D.C.J., Rodriguez-Delgado, C., Scaffidi, A., et al. (2017). Complex sexual deception in an orchid is achieved by co-opting two independent biosynthetic pathways for pollinator attraction. Curr. Biol. 27:1867–1877.

42. Tai, Y., Hou, X., Liu, C., Sun, J., Guo, C., Su, L., et al. (2020). Phytochemical and Comparative Transcriptome Analyses Reveal Different Regulatory Mechanisms in the Terpenoid Biosynthesis Pathways between Matricaria Recutita L. and Chamaemelum Nobile L. BMC genomics 21:169.

43. Lü, P., Yu, S., Zhu, N., Chen, Y. R., Zhou, B., et al. (2018). Genome encode analyses reveal the basis of convergent evolution of fleshy fruit ripening. Nat. Plants 4, 784–791.

44. Li, J., Milne, R. I., Ru, D., Miao, J., Tao, W., et al. (2020). Allopatric divergence and hybridization within Cupressus chengiana (Cupressaceae), a threatened conifer in the northern Hengduan Mountains of western China. Mol. Ecol. 29, 1250–1266.

45. Huang, C.H., Qi, X.P., Chen, D.Y., Qi, J., Ma, H. (2020) Recurrent genome duplication events likely contributed to both the ancient and recent rise of ferns. J. Integr. Plant Biol. 62:433–455.

46. Osuna-Mascaró, C., Rubio de Casas, R., Gomez, J., Loureiro, J., Castro, S., et al. (2022). Hybridization and introgression are prevalent in Southern European Erysimum (Brassicaceae) species. Ann. Bot. https://doi.org/10.1093/aob/mcac048.

47. Young, E., Carey, M., Meharg, A. A., and Meharg, C. (2018). Microbiome and ecotypic adaption of Holcus lanatus (L.) to extremes of its soil pH range, investigated through transcriptome sequencing. Microbiome 6:48.

48. Nevado, B., Atchison, G.W., Hughes, C.E., Filatov, D.A. (2016). Widespread adaptive evolution during repeated evolutionary radiations in New World lupins. Nat. Commun. 7:1–9.

49. Wu, F., Duan, Z., Xu, P., Yan, Q., Meng, M., Cao, M., et al. (2022). Genome and systems biology of Melilotus albus provides insights into coumarins biosynthesis. Plant Biotechnol. J. 20:592–609.

50. Huang, R., Snedden, W., DiCenzo, G. (2022). Reference nodule transcriptomes for Melilotus officinalis and Medicago sativa cv. Algonquin. Grassland Res. 6:e408.

51. Piñeiro-Fernández, L., Byers, K.J., Cai, J., Sedeek, K.E., Kellenberger, R.T., Russo, A., Qi, W., Aquino Fournier, C., Schlüter, P.M. (2019). A phylogenomic analysis of the floral transcriptomes of sexually deceptive and rewarding European orchids, *Ophrys* and *Gymnadenia*. Front. Plant Sci. 10:1553.

52. Peery, R., McAllister, C., Cullingham, C., Mahon, E., Arango-Velez, A., et al. (2021). Comparative genomics of the chitinase gene family in lodgepole and jack pines: contrasting responses to biotic threats and landscape level investigation of genetic differentiation. Botany. 99:355–378.

53. Cai, N., Xu, Y., Chen, S., He, B., Li, G., et al. (2016). Variation in seed and seedling traits and their relations to geo-climatic factors among populations in Yunnan Pine (Pinus yunnanensis). J. For. Res. 27, 1009–1017.

54. Zhao, Z., Luo, Z., Yuan, S., Mei, L., Zhang, D. (2019). Global transcriptome and gene co-expression network analyses on the development of distyly in *Primula oreodoxa*. Heredity. 123:784–794.

55. Pellino M., Hojsgaard D., Schmutzer T., Scholz U., Hörandl E., et al. (2013). Asexual genome evolution in the apomictic Ranunculus auricomus complex: examining the effects of hybridization and mutation accumulation. Mol. Ecol. 22:5908–5921.

56. Yang, Z., Li, W., Su, X., Ge, P., Zhou, Y., et al. (2019) Early Response of Radish to Heat Stress by Strand-Specific Transcriptome and miRNA Analysis. Int. J. Mol. Sci. 20:3321.

57. Zhou, B., Wang, J., Lou, H., Wang, H.Z., Xu, Q.J. (2019a). Comparative transcriptome analysis of dioecious, unisexual floral development in *Ribes diacanthum* pall. Gene 699:43–53.

58. Wickett, N.J., Mirarab, S., Nguyen, N., Warnow, T., Carpenter, E.J., et al. (2014). Phylotranscriptomic analysis of the origin and early diversification of land plants. Proc Natl Acad Sci. 111:4859–68.

59. Meier, S.K., Adams, N., Wolf, M., Balkwill, K., Muasya, A.M., et al. (2018). Comparative RNA-seq analysis of nickel hyperaccumulating and non-accumulating populations of *Senecio coronatus* (Asteraceae). Plant J. 95:1023–1038.

60. Baloun, J., Nevrtalova, V., Kovacova, V., Hudzieczek E., Cegan, R., et al., (2014). Characterization of the HMA7 gene and transcriptomic analysis of candidate genes for copper tolerance in two Silene vulgaris ecotypes. J. Plant Physiol. 171:1188–1196.

61. Clancy, M.V.; Haberer, G.; Jud, W.; Niederbacher, B.; Niederbacher, S.; et al. (2020). Under fire-simultaneous volatilome and transcriptome analysis unravels fine-scale responses of tansy chemotypes to dual herbivore attack. BMC Plant Biol. 20:551.

62. Zhou, T., Luo, X., Yu, C., Zhang, C., Zhang, L., et al (2019b) Transcriptome analyses provide insights into the expression pattern and sequence similarity of several taxol biosynthesis-related genes in three Taxus species. BMC Plant Biol. 19:33.

63. Hodge, B.A., Paul, P.A., Stewart, L.R. (2020). Occurrence and High-Throughput Sequencing of Viruses in Ohio Wheat. Plant Dis. 104:1789–1800.

64. Yu X.L., Wang W.X., Yang H.X., Zhang X.Y., Wang D., et al. (2021b). Transcriptome and comparative chloroplast genome analysis of Vincetoxicum versicolor: insights into molecular evolution and phylogenetic implication. Front. Genet. 12:602528.

65. Lanver, D., Müller, A. N., Happel, P., Schweizer, G., Haas, F. B., et al. (2018). The biotrophic development of Ustilago maydis studied by RNA-seq analysis. Plant Cell 30:300–323.

