## Supplementary material for "Unlocking the hidden genetic diversity of varicosaviruses, the neglected plant rhabdoviruses": Supp Fig. 1

**Supplementary Figure 1**. Stacked bar chart showing the number of previously reported varicosaviruses and in this study.

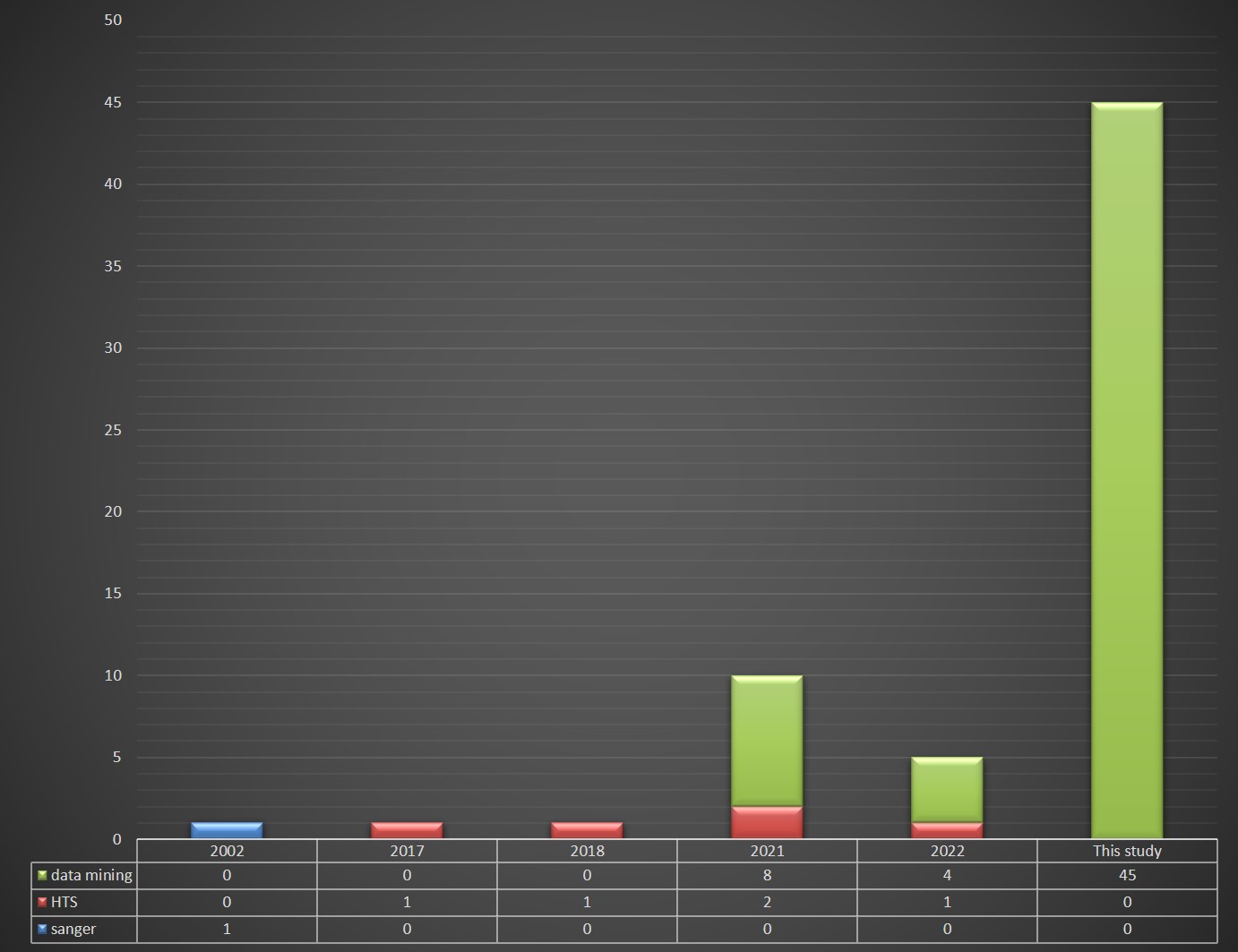
