## Supplementary material for "Unlocking the hidden genetic diversity of varicosaviruses, the neglected plant rhabdoviruses": Supp Table S1

Supplementary Table S1. Virus names, abbreviations and NCBI accession numbers of varicosavirus sequences used in this study

| **Virus name** | **Abbreviation** | **Accession number** |
| --- | --- | --- |
| Allium angulosum virus 1 | AAnV1 | BK059208; BK059209 |
| Alopecurus myosuroides varicosavirus 1 | AMVV1 | LN713933; LN713934 |
| Brassica virus 1 | BrV1 | BK014310; BK014311 |
| lettuce big-vein associated virus | LBVaV | AB075039; AB114138 |
| Lolium virus 1 | LoV1 | BK014312; BK014313 |
| Melampyrum roseum virus 1 | MelRoV1 | BK014314; BK014315 |
| morning glory varicosavirus | MGVV | MW922438; MW922439 |
| Monoclea gottschei varicosavirus | MgVV | OW528612; OW527630 |
| Pinus flexilis virus 1 | PiFleV1 | BK014316 |
| red clover-associated varicosavirus | RCaVV | MF918568; MF918569 |
| Spinach virus 1 | SpV1 | BK061809; BK061810 |
| Tree fern varicosavirus | TfVV | OW528630; OW528632 |
| vitis varicosavirus | VVV | LC604719; LC604720 |
| Xinjiang varicosavirus | XVV | MW897032; MW897033 |
| Zostera-associated varicosavirus 1 | ZaVV1 | BK014484; BK014485 |
